# Exploring the Optimization of Autoencoder Design for Imputing Single-Cell RNA Sequencing Data

**DOI:** 10.1101/2023.02.16.528866

**Authors:** Nan Miles Xi, Jingyi Jessica Li

## Abstract

Autoencoders are the backbones of many imputation methods that aim to relieve the sparsity issue in single-cell RNA sequencing (scRNA-seq) data. The imputation performance of an autoencoder relies on both the neural network architecture and the hyperparameter choice. So far, literature in the single-cell field lacks a formal discussion on how to design the neural network and choose the hyperparameters. Here, we conducted an empirical study to answer this question. Our study used many real and simulated scRNA-seq datasets to examine the impacts of the neural network architecture, the activation function, and the regularization strategy on imputation accuracy and downstream analyses. Our results show that (i) deeper and narrower autoencoders generally lead to better imputation performance; (ii) the sigmoid and tanh activation functions consistently outperform other commonly used functions including ReLU; (iii) regularization improves the accuracy of imputation and downstream cell clustering and DE gene analyses. Notably, our results differ from common practices in the computer vision field regarding the activation function and the regularization strategy. Overall, our study offers practical guidance on how to optimize the autoencoder design for scRNA-seq data imputation.

## Introduction

Single-cell RNA-sequencing (scRNA-seq) enables the measurement of genome-wide gene expression at the single-cell level ^1–3^. State-of-the-art scRNA-seq technologies can measure tens of thousands of genes and up to millions of cells ^4^, allowing the investigation of cell-to-cell heterogeneity ^5^, the identification of cell types ^6^, and the inference of cell state transitions ^7^. One characteristic of scRNA-seq data is the high proportion of zeros (i.e., high sparsity). Depending on the sequencing platform and sequencing depth, the zero proportion can range from 50% to more than 90% ^8^. Two types of zeros exist in scRNA-seq data: biological zeros and non-biological zeros ^9^. Biological zeros indicate the actual absence of gene expression in cells, while non-biological zeros originate from technical biases and noise in scRNA-seq experiments ^10^. Without spike-ins or prior biological knowledge, it is difficult to distinguish between these two types of zeros in scRNA-seq data ^8^.

The high sparsity of scRNA-seq data hinders data analysis. Many computational methods have been developed to reduce data sparsity, and they can be divided into three broad categories ^8^. First, model-based imputation methods use probabilistic models to describe gene expression distributions in scRNA-seq data (e.g., scImpute ^11^ and BISCUIT ^12^). These methods aim to first distinguish between biological zeros and non-biological zeros and then only impute the identified non-biological zeros. Second, data-smoothing methods modify each cells’ gene expression levels based on similar cells’ gene expression levels (e.g., MAGIC ^13^ and DrImpute ^14^). Two cells may be defined as similar if they share common neighboring cells in a low-dimensional space. Unlike model-based imputation methods, data-smoothing methods do not distinguish non-biological zeros and would alter all gene expression levels, including the nonzeros. Third, data-reconstruction methods first learn a latent low-dimensional space of cells and then reconstruct cells in the original high-dimensional space (e.g., ZIFA ^15^ and DCA ^16^). Most data-reconstruction methods use the reconstructed data to replace all zeros in the original data but keep the nonzeros unchanged.

Among the data-reconstruction methods, autoencoder-based methods are popular for their imputation accuracy and scalability ^17,18^. Specifically, an autoencoder is a neural network that contains an encoder for dimension reduction and a decoder for data reconstruction. Then the reconstructed data are regarded as the imputed scRNA-seq data. Despite the wide use of autoencoders, the scRNA-seq field lacks a formal discussion on how to optimize the autoencoder design, including the neural network architecture, the activation function, and the parameter regularization ^8^. Existing autoencoder-based methods either adopt the wisdom in other fields (e.g., computer vision) or design autoencoders with limited justification.

Here, we conduct an empirical study to explore how to optimize the autoencoder design for imputing scRNA-seq data. Our study consists of three investigations. The first investigation is the optimization of autoencoder design for imputation accuracy. In detail, we generate 36 semi-synthetic scRNA-seq datasets with artificial zeros (whose original, non-missing values are known) by applying three masking schemes to 12 real scRNA-seq datasets. Then, we train autoencoders with varying depths (depth means the number of layers) and widths (width means the number of nodes per layer), seven activation functions, and two regularization strategies on each semi-synthetic dataset. Next, we evaluate the trained autoencoders’ imputation accuracies in terms of the normalized root mean squared error (NRMSE) and the Pearson correlation coefficient, which are calculated between the imputed and original values of the artificial zeros. From the evaluation results, we find the autoencoder designs that lead to the best imputation accuracies.

The second investigation is the optimization of autoencoder design for cell clustering. In detail, we train autoencoders with the aforementioned designs on 20 real scRNA-seq datasets containing curated cell type information. Next, to examine the impact of autoencoder design on downstream cell clustering, we apply the trained autoencoders for imputation and evaluate the clustering accuracies on the imputed datasets in terms of the adjusted rand index (ARI) and the adjusted mutual information (AMI).

The third investigation is the optimization of autoencoder design for differentially expressed (DE) gene identification. In detail, we simulate 20 synthetic datasets with ground-truth DE genes by applying the scDesign simulator ^19^, which is trained on 20 real scRNA-seq datasets (see Methods). Then we train autoencoders with the aforementioned designs on these synthetic datasets. Next, to examine the impact of autoencoder design on downstream DE gene analysis, we apply the trained autoencoders for imputation and evaluate the DE gene identification accuracies on the imputed datasets in terms of the precision, recall, and true negative rate (TNR).

In the above three investigations, we consider many neural network depths, widths, activation functions, and regularization strategies. Since a full combination of those design aspects is computationally infeasible, we adopt a sequential, greedy search strategy to explore the best design. In each investigation, we first examine a full combination of neural network depths and widths, the two most important aspects, with the other design aspects fixed as in common practices (see Results). Based on the optimized combination of width and depth, we next optimize the activation function. Finally, given the optimized width, depth, and activation function, we optimize the regularization strategy.

Our results yield the following guidelines about autoencoder design for scRNA-seq data imputation (Table 1). First, deeper and narrower autoencoders lead to more accurate imputation, cell clustering, and DE gene identification, yet the benefit of depth saturates at 10 hidden layers. Second, the sigmoid and tanh activation functions consistently have the best performance in all evaluations. Third, parameter regularization is critical to the performance of autoencoder-based imputation methods. In particular, weight decay regularization is more capable of improving cell clustering and DE gene analysis, while dropout regularization shows superiority in improving the overall imputation accuracy for the artificial zeros. Moreover, the optimal degree of regularization is dataset-specific.

**Table 1.** Guidelines of autoencoder design for scRNA-seq data imputation. Each row represents one of four design characteristics. Each column represents one of three evaluation metrics. Each entry corresponds to the marginally optimized design characteristic under each evaluation metric.

| Autoencoder design characteristics | Evaluation metrics |  |  |
| --- | --- | --- | --- |
|  | Imputation accuracy | Cell clustering | DE gene identification |
| Number of hidden layers | $\geq 10$ | $\geq 10$ | $\geq 5$ |
| Number of units per hidden layer | 32 | Robust | Robust |
| Activation function | Sigmoid or Tanh | Sigmoid or Tanh | Sigmoid or Tanh |
| Regularization strategy | Dropout | Weight Decay | Weight Decay |

Our findings suggest that many autoencoder-based imputation methods have used suboptimal autoencoder designs, including shallower and wider autoencoders and the ReLU activation function. Moreover, our findings highlight the importance of using empirical benchmarking to optimize autoencoder designs, and more generally, neural network designs, in bioinformatics research.

## Results

### Impacts of autoencoder architecture (depth and width) on imputation accuracy

We collect 12 real scRNA-seq datasets to evaluate the overall imputation accuracy of a variety of autoencoder architectures. These datasets cover a wide range of cell types, sequencing depths, zero proportions, and experimental platforms (Supplementary Table S1). We apply three masking schemes (i.e., random masking, double exponential masking, and medium masking, which reflect different degrees of dependence of missingness on the actual values; see Methods) to these 12 real datasets, obtaining 36 masked datasets. To evaluate the imputation accuracy on each masked dataset, we calculate the NRMSE and the Pearson correlation coefficient between the masked values and the imputed values (Methods), referred to as the “imputation NRMSE” and the “imputation correlation,” respectively, in the following text.

We build autoencoders of various architectures by increasing the depth (i.e., the number of hidden layers) from 1 to 15. For each depth, we set the width (i.e., the number of hidden units per layer) to 32, 64, 128, or 256. All hidden layers are fully connected with the same width. In total, we have 60 autoencoder architectures, corresponding to 15×4 depth-width combinations. We choose the rectified linear unit (ReLU) function ^20^ as the activation function and train the autoencoders by the Adam optimization algorithm ^21^ (Methods). For each autoencoder architecture and each masked dataset, we set 10 random seeds in the autoencoder training to obtain 10 autoencoders, whose imputation NRMSEs and imputation correlations are averaged to represent the overall imputation accuracy of the autoencoder architecture on the masked dataset.

**Figure 1.**
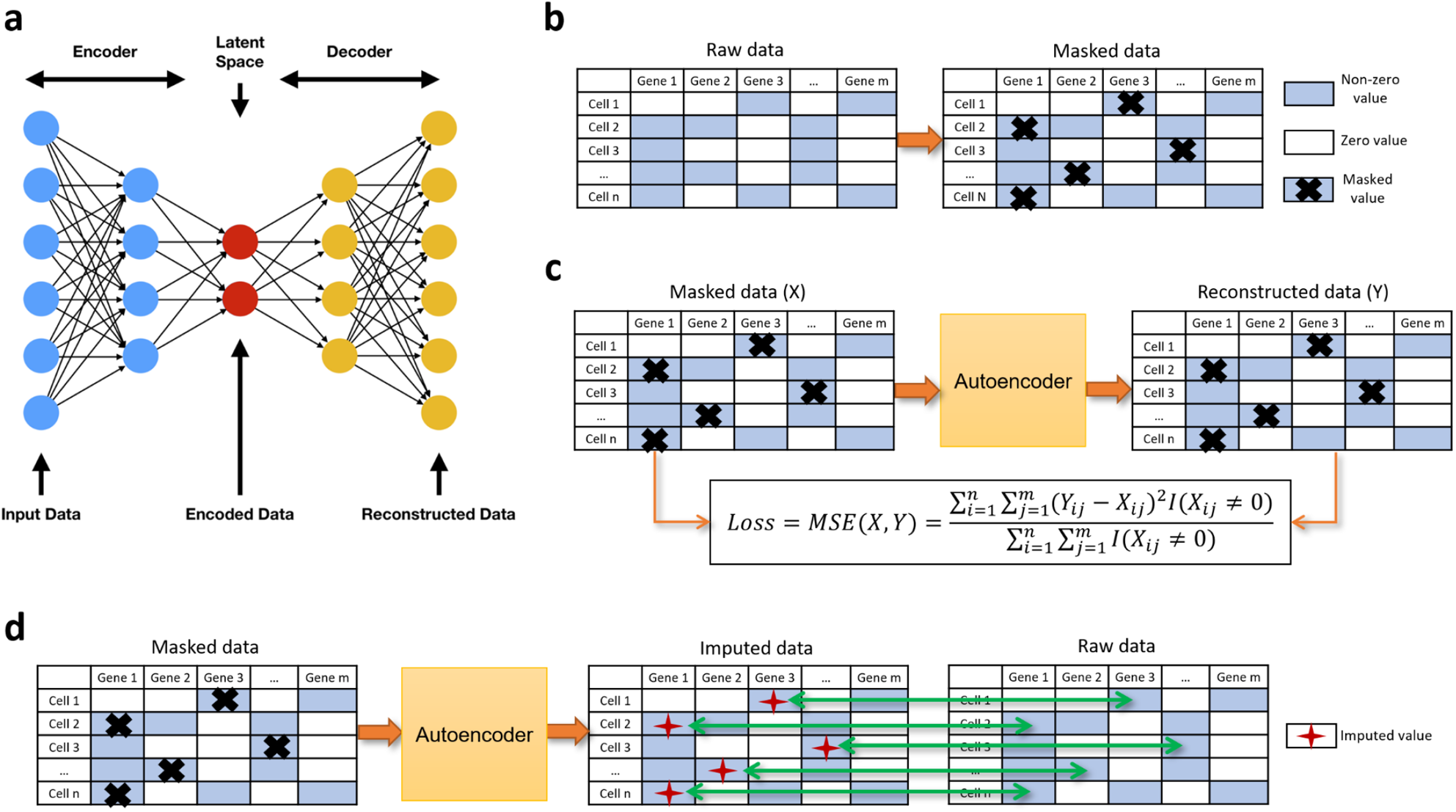
Autoencoder and the measurement of imputation accuracy. **a**, The basic structure of an autoencoder. **b**, The introduction of artificial zeros by masking. **c**, The training of an autoencoder for imputation. The reconstructed data are the autoencoder’s output during the training process. **d**, The calculation of an autoencoder’s imputation accuracy on masked values. The imputed data are the final autoencoder’s output after the training process.

Under the random masking scheme, Figure 2a–b show the impacts of autoencoder depth-width combinations on the imputation NRMSE and the imputation correlation. First, deeper autoencoders achieve lower imputation NRMSEs and higher imputation correlations. The benefit of depth is more prominent when the autoencoder has no more than 10 layers. Second, narrower autoencoders (with 32 hidden units per layer) typically have higher imputation accuracy than wider autoencoders (with 64 or more hidden units per layer) of the same depth. This finding is consistent with the observation that deeper and narrower neural networks perform better in computer vision tasks (e.g., image classification and object detection) ^22,23^. Under the double exponential masking scheme, we observe a similar relationship between the autoencoder depth-width combinations and the imputation accuracy, except for the dataset `bmmc` (Supplementary Figure S1). However, the relationship no longer holds under the median masking scheme: while deeper autoencoders still have better imputation accuracy on certain datasets, the width does not have a significant impact on the imputation accuracy (Supplementary Figure S2). Among the three masking schemes, random masking has the highest imputation accuracy, followed by double exponential masking and then median masking. Note that under median masking, the imputation accuracy is low on most datasets, indicated by many larger-than-one imputation NRMSEs and close-to-zero imputation correlations (Supplementary Figure S2).

**Figure 2.**
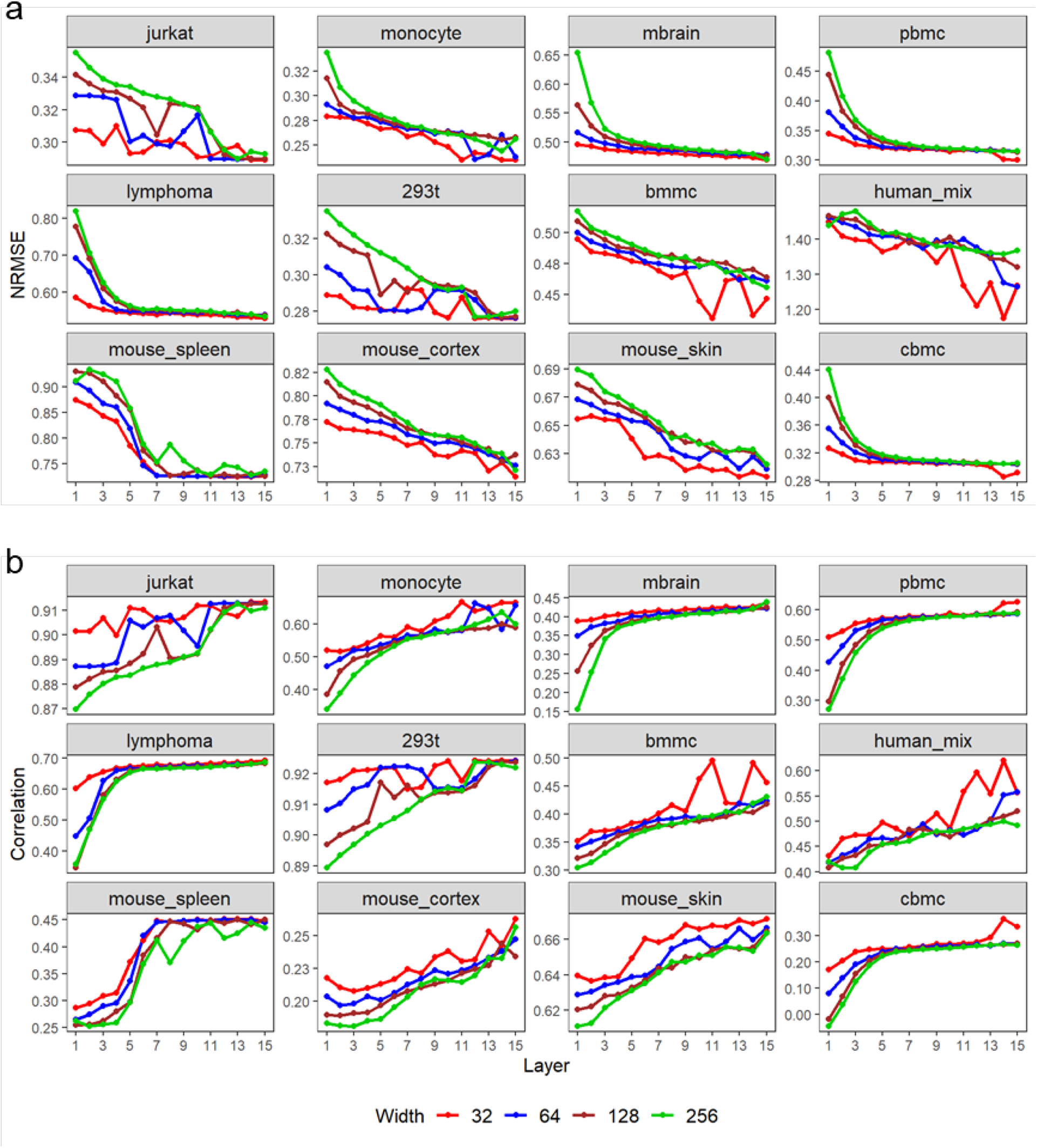
The impacts of autoencoder depth and width on the imputation NRMSE (**a**) and the imputation correlation (**b**) based on the random masking scheme. Each point is the average of the results obtained from 10 random seeds used for autoencoder training.

A possible explanation of the results is that, among the three masking schemes, random masking best preserves gene-gene correlations. Since random masking is performed independent of gene expression levels, the gene-gene correlations in the masked data are unbiased estimates of the gene-gene correlations in the original, unmasked data. Unlike random masking, double exponential masking assumes an inverse relationship between a gene’s masking probability and average expression (i.e., the probability of masking increases as a gene’s average expression level decreases); hence, the gene-gene correlations in the masked data are no longer unbiased estimates of the gene-gene correlations in the original data. While random masking and double exponential masking are both stochastic schemes, median masking is a deterministic scheme where the 50% lowest expression levels of all genes are masked as zeros. Hence, median masking distorts gene-gene correlations to the greatest extent among the three masking schemes. Under the hypothesis that gene-gene correlations play a crucial role in determining the imputation accuracy of an autoencoder, it is unsurprising that the best imputation accuracy is achieved under the random masking scheme, followed by the double exponential masking scheme. Hence, the following analysis on imputation accuracy will focus on random masking and double exponential masking.

### Impacts of activation function on imputation accuracy

An activation function is a nonlinear transformation applied to generate the hidden units in an autoencoder ^24^. It provides an autoencoder with the capacity to learn complex nonlinear relationships among features, e.g., nonlinear interactions among genes in scRNA-seq data. ReLU is a widely used activation function in autoencoder-based imputation methods, motivated by its success in computer vision ^24^. However, justification is lacking for using ReLU to impute scRNA-seq data, and no empirical comparison has been done between ReLU and other activation functions.

Here, we train autoencoders with seven activation functions, including sigmoid, tanh, ReLU, LeakyReLU (with two hyperparameter settings) ^25^, ELU ^26^, and SELU ^27^, to evaluate the impacts of activation functions on the imputation accuracy (Methods). For each activation function, we impute 24 masked datasets (the aforementioned 12 scRNA-seq datasets after random masking or double exponential masking) by training 20 autoencoders using 20 random seeds on each masked dataset. Figure 3 and Supplementary Figure S3 compare the seven activation functions’ resulting imputation NRMSEs and imputation correlations under the random masking and double exponential masking schemes. Surprisingly, the most popular ReLU function is not the top performer. Instead, the sigmoid and tanh functions outperform the other activation functions on all datasets under both masking schemes. Additionally, the sigmoid and tanh functions result in significantly smaller variances of the imputation NRMSEs and the imputation correlations than the other functions do, indicating that the sigmoid and tanh functions lead to more stable imputation accuracy. Between sigmoid and tanh, although they have similar performance, sigmoid has higher imputation accuracy on the datasets `pbmc` and `human_mix` and more stable imputation accuracy on the datasets `mbrain`, `pbmc`, `human_mix`, and `mouse_cortex`.

**Figure 3.**
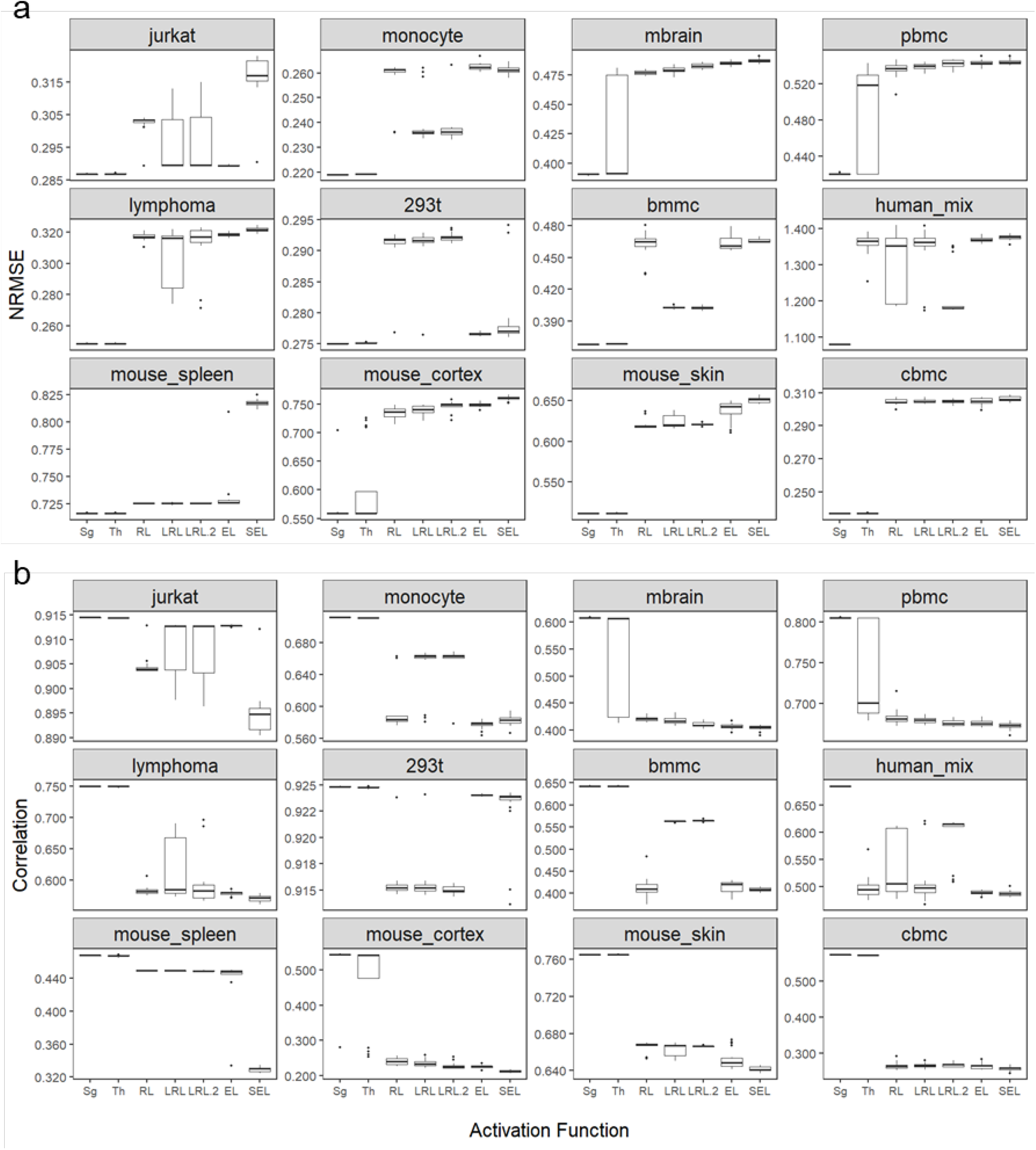
The impact of activation functions on the imputation NRMSE (**a**) and the imputation correlation (**b**) based on the random masking scheme. Sg: sigmoid; Th: tanh; RL: ReLU; LRL: LeakyReLU (*α* = 0.01); LRL.2: LeakyReLU (*α* = 0.2); EL: ELU; SEL: SELU. Each boxplot shows the results obtained from 20 random seeds used for autoencoder training.

The comparison of the seven activation functions reveals three insights. First, unlike ReLU that introduces sparsity into hidden layers (i.e., many hidden units are zeros) ^25^, sigmoid and tanh generate dense hidden layers (i.e., many hidden units are nonzeros). Our empirical results suggest that dense hidden layers lead to higher imputation accuracy for scRNA-seq data. Second, Leaky ReLU, ELU, and SELU are modified forms of ReLU that output a close-to-zero negative value when the input is negative (Methods). They generate pseudo-sparsity in the hidden layers to avoid the “dead ReLU” problem (i.e., most hidden units are zeros, causing the autoencoder to stop learning) ^28^. However, our empirical results show no consistent improvement of these modified ReLU functions over ReLU in terms of imputation accuracy, though these modified ReLU functions lead to dense hidden layers as sigmoid and tanh do. A possible interpretation is that sigmoid and tanh have continuous derivatives, while the modified ReLU functions have discontinuous derivatives. Third, we do not observe vanishing gradients or exploding gradients ^29^, two common problems in training deep neural networks using sigmoid or tanh. We hypothesize that appropriate preprocessing and normalization of scRNA-seq data help stabilize gradients in the training process.

### Impacts of regularization on imputation accuracy

Some autoencoder-based imputation methods use weight decay or dropout as the regularization strategy ^30,31^. However, the selection of the regularization strategy and the corresponding hyperparameters (i.e., *λ* in weight decay and *p* in dropout, Methods) are mostly ad hoc ^8^. To examine the impact of regularization on imputation accuracy, we train autoencoders with weight decay or dropout as the regularization strategy (Methods) to impute the aforementioned 24 masked datasets (the 12 real scRNA-seq datasets after random masking or double exponential masking). We vary *λ* ∈ [1e-7, 5e-4] and *p* ∈ [0.01, 0.4] and compare the resulting imputation NRMSEs and imputation correlations (Method; Figure 4 and 5; Supplementary Figure S4 and S5). All autoencoders have the same architecture and activation function: 10 fully connected hidden layers, 32 hidden units per hidden layer, and the sigmoid activation function. For each dataset and regularization strategy (with hyperparameter), we use 10 random seeds to train 10 autoencoders and average the resulting imputation NRMSEs and imputation correlations.

**Figure 4.**
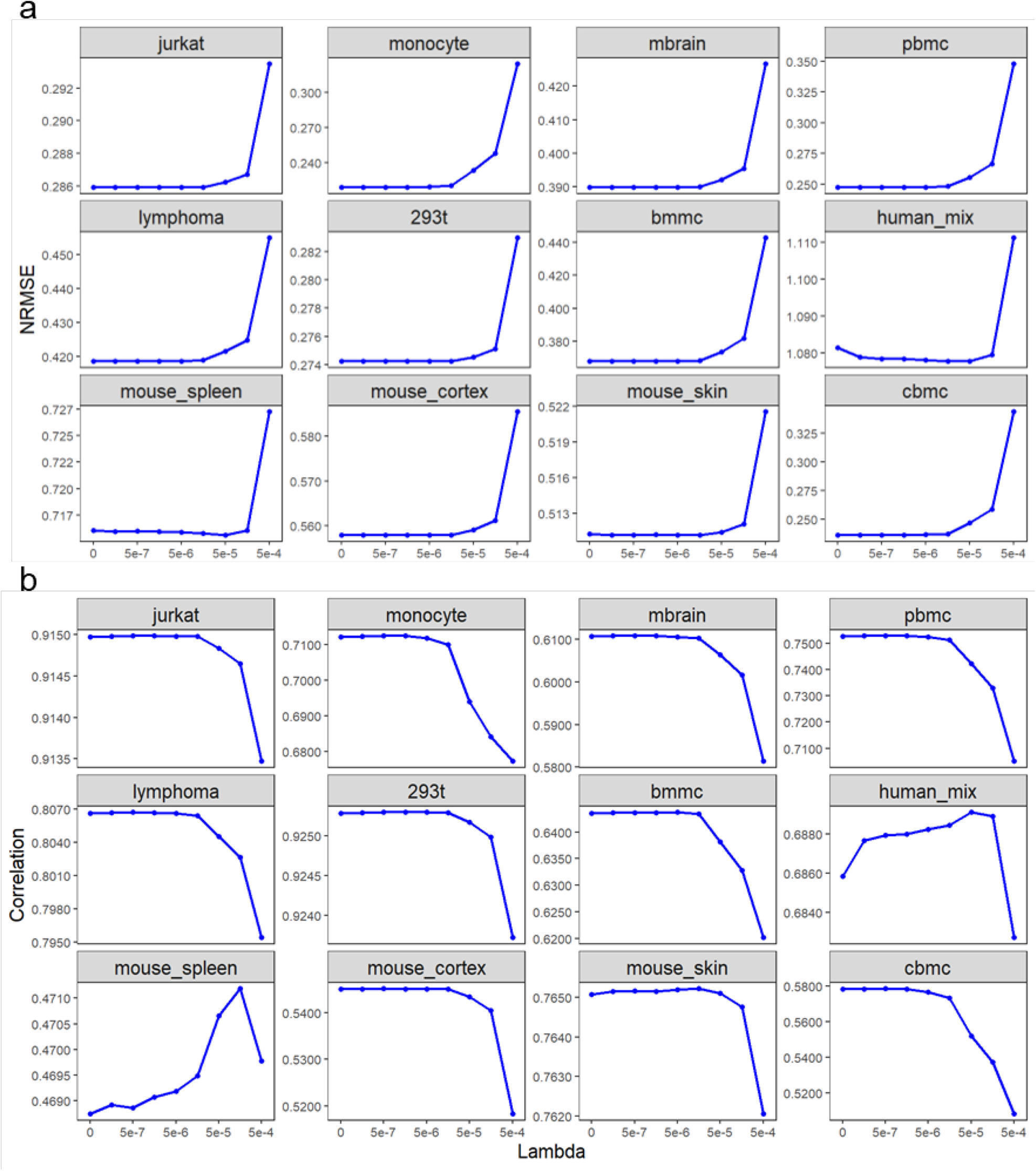
The impact of weight decay regularization on the imputation NRMSE (**a**) and the imputation correlation (**b**) based on the random masking scheme. Each point is the average of the results obtained from 10 random seeds used for autoencoder training.

**Figure 5.**
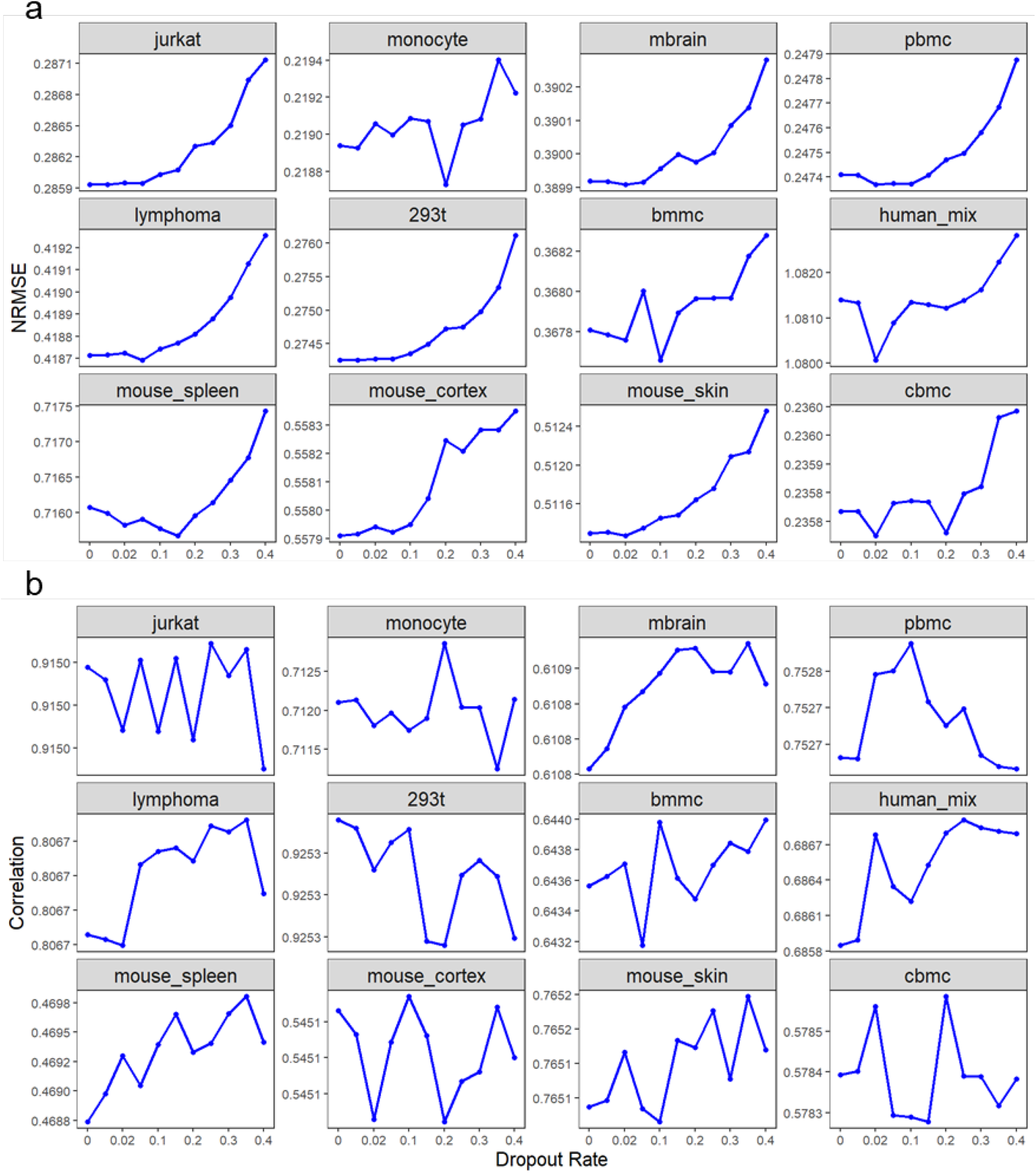
The impact of dropout regularization on the imputation NRMSE (**a**) and the imputation correlation (**b**) based on the random masking scheme. Each point is the average of the results obtained from 10 random seeds used for autoencoder training. Some labels on y-axis are identical due to the low accuracy variability under different dropout rates.

Under random masking, the weight decay strategy barely improves the imputation accuracy except on datasets `mouse_spleen` and `human_mix`. Larger values of *λ* even lead to worse imputation accuracy, suggesting over-regularization (Figure 4). In contrast, the dropout strategy with a proper dropout rate *p* improves the imputation NRMSEs on six datasets and the imputation correlations on 11 datasets. The optimal range of *p* depends on the evaluation metric: 0.02–0.2 for imputation NRMSE and 0.2–0.4 for imputation correlation (Figure 5). Hence, *p* = 0.2 is a reasonable choice.

Under double exponential masking, both the weight decay and dropout strategies improve the imputation accuracy (Supplementary Figure S4 and S5). Specifically, the weight decay strategy reduces the imputation NRMSEs on six datasets and increases the imputation correlations on 11 datasets (Supplementary Figure S4). The optimal range of *λ* depends on the evaluation metric: 1e-7– 5e-5 for imputation NRMSE and 5e-6–5e-4 for imputation correlation. Hence, 5e-6–5e-5 is a reasonable range for *λ*. The dropout strategy exhibits a stronger positive impact on the imputation accuracy, with the imputation NRMSEs reduced on 10 datasets and the imputation correlations increased on all 12 datasets (Supplementary Figure S5). The optimal range of the dropout rate *p* is similar to that under random masking.

We summarize the impact of regularization on the imputation accuracy from three perspectives. First, compared with the weight decay strategy, the dropout strategy is more capable of improving both the imputation NRMSE and the imputation correlation under both random masking and double exponential masking. Second, both the weight decay and dropout strategies are more effective under double exponential masking than random masking. As discussed in “Impacts of autoencoder architecture (depth and width) on imputation accuracy,” double exponential masking masks small nonzero values in the scRNA-seq data matrix, thus distorting gene-gene correlations. In contrast, random masking does not have this issue. Hence, we hypothesize that regularization has a bigger benefit under double exponential masking because regularization’s increase in robustness is more needed after double exponential masking’s distortion on gene-gene correlations. Third, the optimal hyperparameters of regularizations largely depend on the datasets and masking schemes, making it impossible to find a universally optimized hyperparameter setting. Interestingly, compared with the imputation NRMSE, the imputed correlation requires a higher degree of regularization to achieve its optimum, under both the weight decay and dropout strategies. Finding the reason underlying this phenomenon requires future research.

### Impacts of autoencoder design (architecture, activation function, and regularization) on cell clustering

The ultimate goal of imputation is to improve the downstream analysis through the enhancement of signals in the sparse scRNA-seq data ^32^. We collect 20 real scRNA-seq datasets containing annotated cell types to examine the impact of autoencoder design on cell clustering (Supplementary Table S2). Here, the datasets are different from those used in the evaluation of the imputation accuracy because evaluating clustering accuracy does not require the knowledge of non-missing values. Specifically, we first conduct K-means clustering on each original dataset and calculate the adjusted rand index (ARI) and the adjusted mutual information (AMI) to measure the clustering performance (Methods). Note that we choose K-means clustering instead of the more popular graph-based Louvain or Leiden clustering in the Seurat package ^33^ because we want to set K, the number of clusters, to the number of cell types for fair evaluation. While setting K to a specific number is natural for K-means clustering, it requires manual tuning for Louvain and Leiden clustering and is too labor-intensive for our evaluation. Second, we train autoencoders with various architectures, activation functions, and regularization strategies to impute the 20 datasets. Finally, we conduct K-means clustering on each imputed dataset and calculate the corresponding ARI and AMI.

Figure 6a and Supplementary Figure S6a show the impact of autoencoder architecture on cell clustering. Similar to the previous analysis, we increase the depth of autoencoders from 1 to 15 and set the width to 32, 64, 128, and 256 for each depth, respectively. All hidden layers in each autoencoder are fully connected with the same width. In total, we have 60 (15×4) depth-width combinations. We use the sigmoid activation function because of its superior performance in the previous imputation accuracy evaluation. On each dataset, we use five random seeds for autoencoder training to obtain five imputed datasets, on which we calculate the imputation ARIs and the imputation AMIs; then we report the average imputation ARI and the average imputation AMI. Our results, which are consistent between ARIs and AMIs, show that deeper autoencoders lead to a larger improvement on cell clustering accuracy than their shallower counterparts do. The benefit of depth saturates after 10 hidden layers. Unlike the autoencoder depth, the autoencoder width has no obvious impact on cell clustering accuracy (Figure 6a; Supplementary Figure S6a). Surprisingly, imputation does not always improve cell clustering accuracy: imputation only improves ARIs on eight datasets and AMIs on four datasets, regardless of autoencoder architectures.

**Figure 6.**
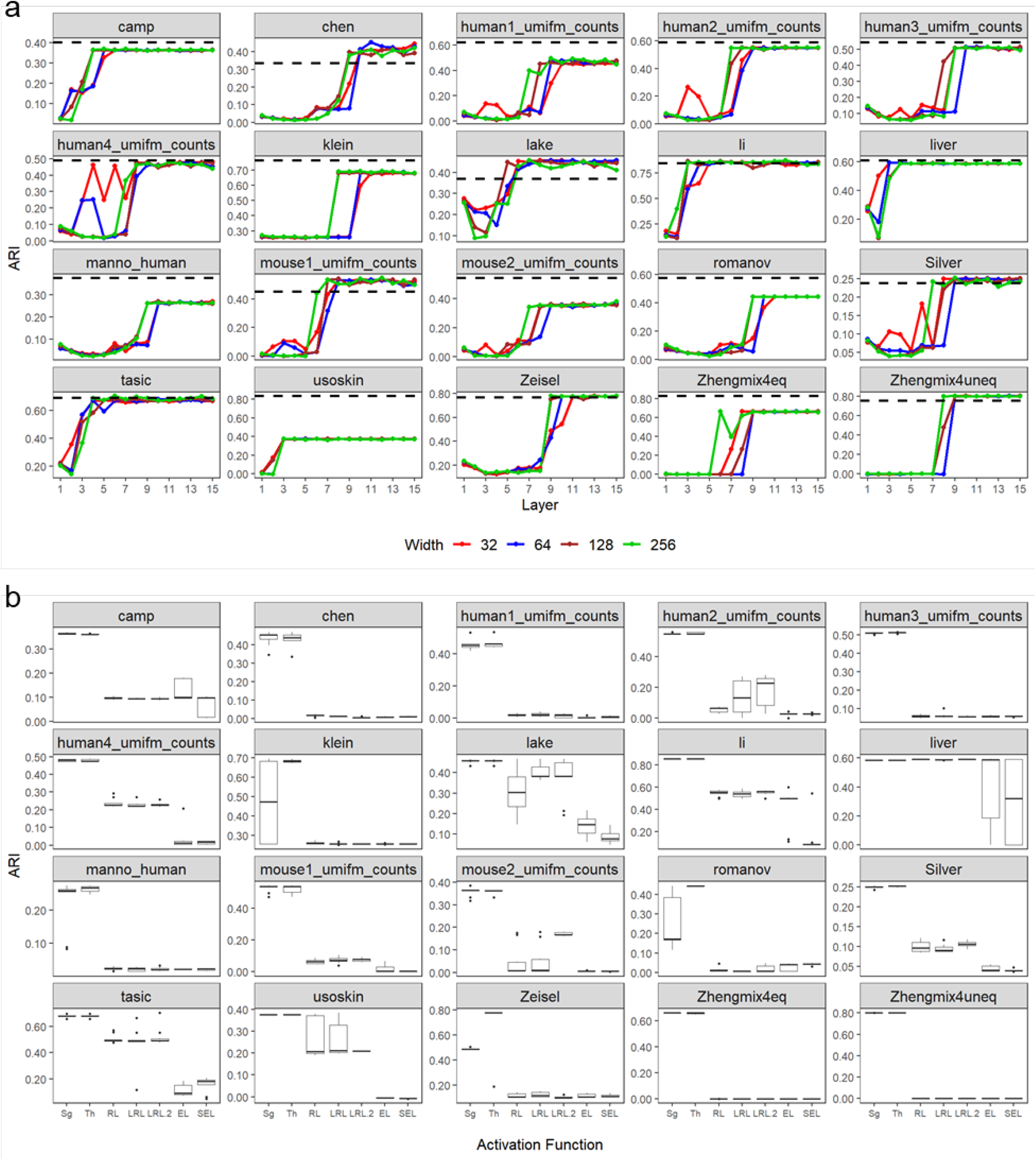
The impact of autoencoder design on cell clustering accuracy measured by ARI. **a,** Depth and width. **b**, Activation function. The dash lines in (**a**) show the cell clustering performance without imputation. Each point in (**a**) is the average of the results obtained from five random seeds used for autoencoder training. Each boxplot in (**b**) shows the results obtained from 10 random seeds used for autoencoder training.

Figure 6b and Supplementary Figure S6b show the impact of activation functions on cell clustering. Similar to the previous analysis, we train autoencoders with seven activation functions, including sigmoid, tanh, ReLU, LeakyReLU (with two different hyperparameters), ELU, and SELU. For each activation function, we impute the 20 scRNA-seq datasets using 10 autoencoders trained under 10 random seeds. All autoencoders have 10 fully connected hidden layers with 32 hidden units per layer, an optimized architecture based on our previous analysis. We observe that sigmoid and tanh outperform other activation functions on all datasets in terms of both imputation AMI and imputation ARI. They also lead to more stable cell clustering accuracy than other activation functions do. The performance of sigmoid and tanh is similar except for datasets `mouse_cortex` and `Klein`, where tanh has a slight advantage over sigmoid.

Figure 7 and Supplementary Figure S7 show the impact of regularization on cell clustering. Similar to the previous analysis, we use weight decay and dropout strategies and vary *λ* ∈ [1e-7, 5] and *p* ∈ [0.01, 0.4] (Methods). Based on our previous analysis, all autoencoders have 10 fully connected hidden layers, with 32 hidden units per layer, and the sigmoid activation function. Interestingly, the weight decay strategy significantly improves cell clustering accuracy—with the optimized weight decay hyperparameter *λ*’s (mostly between 0.01 and 0.1), all 20 datasets have improved ARIs, and 18 datasets have improved AMIs. However, the dropout strategy does not lead to same improvement— with the optimized dropout hyperparameter *p*’s (in a broad range), only eight datasets have improved ARIs, and only four datasets have improved AMIs. Hence, unlike the previous imputation accuracy evaluation, which prefers the dropout strategy, here the weight decay strategy is preferred. However, we do not want to over-interpret the results because the autoencoders we use do not consistently lead to better ARIs and AMIs, suggesting that imputation might not be needed for cell clustering, consistent with our previous report ^9^.

**Figure 7.**
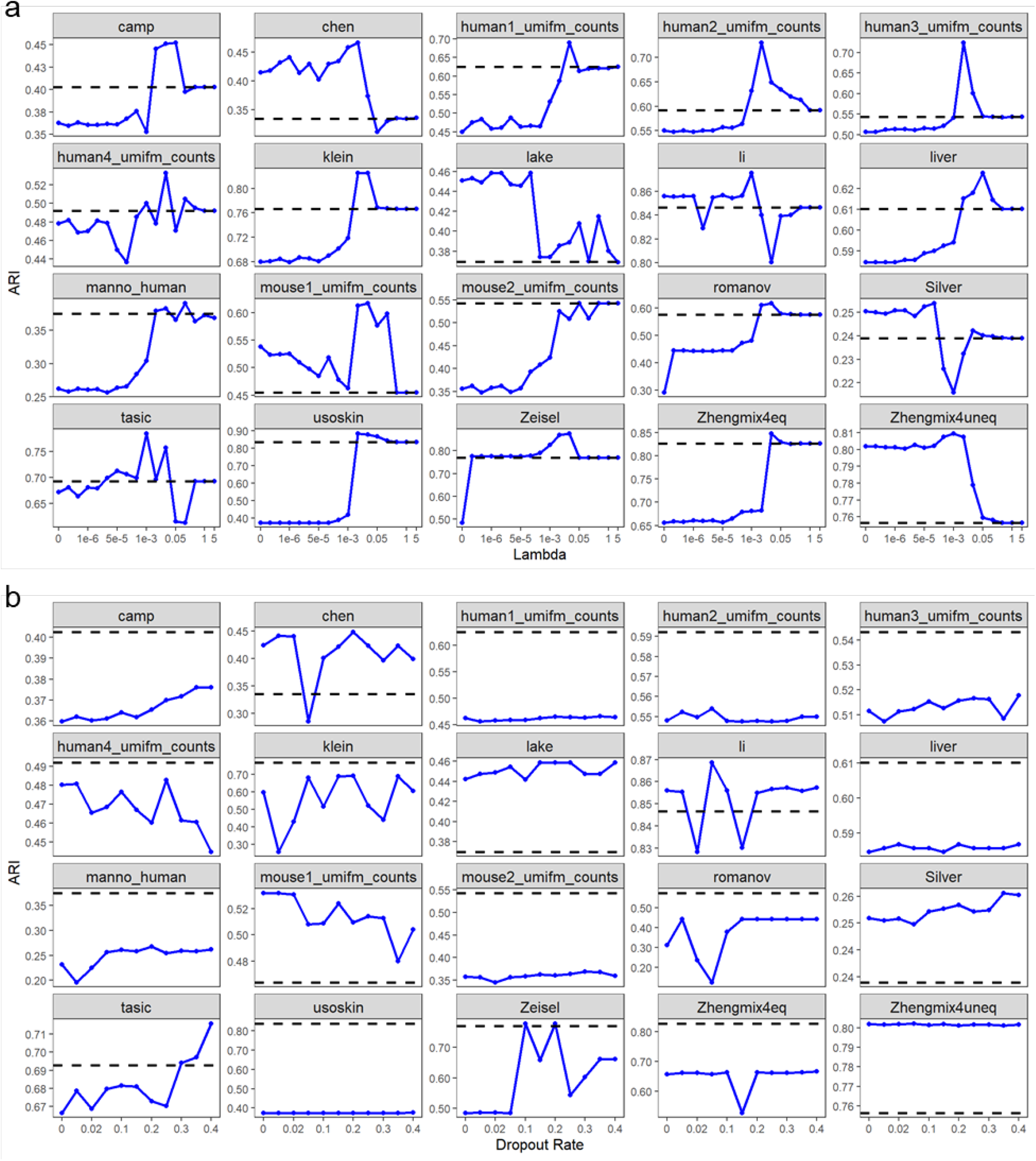
The impact of autoencoder regularization on cell clustering accuracy measured by ARI. **a**, Weight decay regularization. **b**, Dropout regularization. The dash lines show the cell clustering performance without imputation. Each point is the average of the results obtained from five random seeds used for autoencoder training.

### Impact of autoencoder design (architecture, activation function, and regularization) on DE gene analysis

The signal enhancement in scRNA-seq data by imputation is expected to benefit another important downstream analysis—the identification of differentially expressed (DE) genes. To examine the impact of autoencoder design on DE gene analysis, we utilize the simulator scDesign ^19^ to generate 20 synthetic datasets with ground-truth DE genes (Methods). Each synthetic dataset is generated by learning the distributions of genes’ expression levels in one real scRNA-seq dataset (20 real datasets in total; Supplementary Table S3). These real datasets (and their synthetic counterparts) cover a wide range of biological and technical conditions. We use synthetic data in this analysis since the ground-truth DE genes are unknown in real scRNA-seq datasets.

After simulation, we apply the MAST method ^34^ to the pre-imputed synthetic datasets to identify DE genes and calculate the corresponding precision, recall, and true negative rate (TNR), which are considered as the baseline accuracies. Next, to impute each synthetic dataset, we train autoencoders with various architectures, activation functions, and regularization strategies. Finally, we apply MAST to each imputed dataset and calculate the corresponding precision, recall, and TNR (Methods).

Figure 8a, Supplementary Figure S8a, and S10a show the impact of autoencoder architecture on the recall, precision, and TNR. The settings of depth, width and activation function for autoencoders are the same as in the evaluation of cell clustering (1 to 15 fully-connected hidden layers; 32, 64, 128, or 256 hidden units per layer; sigmoid activation function). On each synthetic dataset, we set five random seeds in the autoencoder training to obtain five imputed datasets, whose precisions, recalls, and TNRs are averaged to represent the autoencoder’s performance. First, deeper autoencoders lead to better recalls on 10 datasets, while the benefit of depth saturates after five layers (Figure 8a). In contrast, width has no significant improvement on the recall. Unexpectedly, 11 synthetic datasets show no improvement in recall after imputation, regardless of the autoencoder architecture. Second, depth and width have no significant impact on the precision, except for the datasets `Interneurons`, `Epithelial_cells`, and `astrocytes`, where deeper autoencoders lead to slightly higher precision (Supplementary Figure S8a). Overall, 19 synthetic datasets have improved precision after imputation. Third, the impact of depth and width on the TNR is limited, because all TNRs are already close to one before imputation, and most TNRs are increased by less than 0.05 after imputation (Supplementary Figure S10a). Altogether, our results suggest that autoencoder-based imputation benefits the precision instead of the recall or the TNR, making the identification of DE genes more conservative.

**Figure 8.**
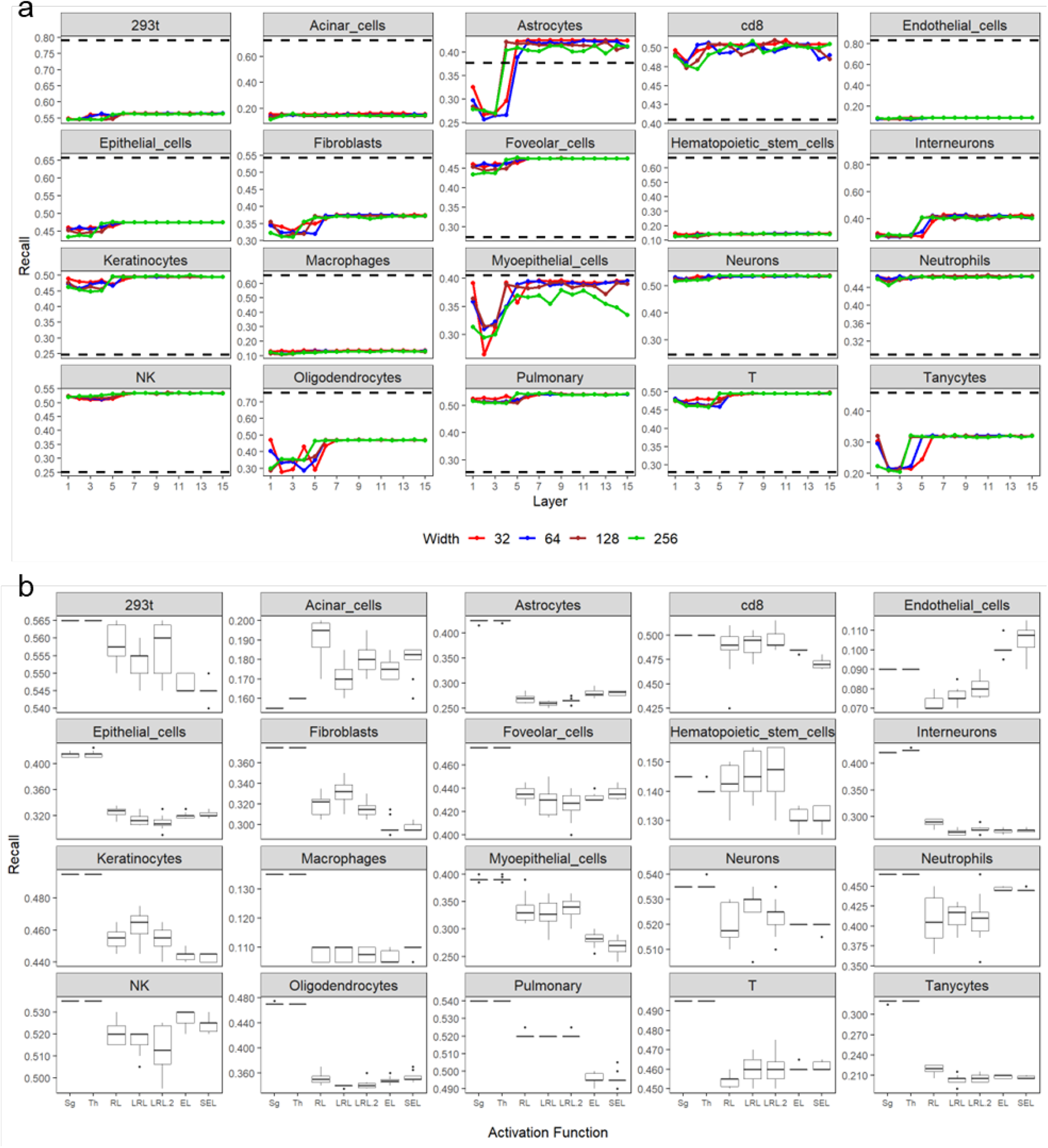
The impact of autoencoder design on DE gene identification accuracy measured by recall. **a,** Depth and width. **b**, Activation function. The dash lines in (**a**) show the recall without imputation. Each point in (**a**) is the average of the results obtained from five random seeds used for autoencoder training. Each boxplot in (**b**) shows the results obtained from 10 random seeds used for autoencoder training.

Figure 8b, Supplementary Figure S8b, and S10b show the impact of activation functions on the recall, precision, and TNR. Again, we train autoencoders with seven activation functions; for each function, we impute the aforementioned 20 synthetic datasets using 10 autoencoders trained by setting 10 random seeds. All autoencoders have 10 fully connected hidden layers with 32 hidden units per layer. In terms of the recall, sigmoid and tanh outperform the other activation functions on 15 datasets (Figure 8b). The advantage of sigmoid and tanh is similar for the precision—they outperform the other activation functions on 12 datasets (Supplementary Figure S8b). All activation functions have similar performance based on the TNR, which is consistently close to one (Supplementary Figure S10b). Overall, sigmoid and tanh have similar performance, and they both provide a more stable improvement in the identification of DE genes than other activation functions do.

Figure 9, Supplementary Figure S9, and S11 show the impact of regularization on the recall, precision, and TNR. Again, we add either weight decay or dropout to autoencoders and adjust the corresponding hyperparameter, as in the previous analysis. All autoencoders have 10 fully connected hidden layers, with 32 hidden units per layer, and the sigmoid activation function. We observe that, compared to dropout, weight decay exhibits stronger improvement in recall, precision, and TNR: weight decay outperforms the no-regularization autoencoders on 20, 12, and 20 datasets, respectively, with the optimal hyperparameter *λ*’s (in the range of [5e-6, 0.05]) (Figure 9a, Supplementary Figure S9a, and S11a); in contrast, dropout outperforms the no-regularization autoencoders on 17, 6, and 17 datasets, respectively, with the optimal *p*’s (in the range of [0.05, 0.25]) (Figure 9b, Supplementary Figure S9b, and S11b). Moreover, weight decay with the optimal *λ*’s improves the recall, precision, and TNR from the baseline values (before imputation) on every dataset, while dropout with the optimal *p*’s fails to improve the recall and TNR from the baseline values on 6 and 20 datasets, respectively. The preference of weight decay over dropout is similar to our evaluation of cell clustering but different from our evaluation of imputation accuracy. A possible reason is that we use manual masking (random or double exponential) in the evaluation of imputation accuracy but not in the evaluation of cell clustering or DE gene identification. Our results reveal the importance of evaluation metrics in autoencoder design. However, despite the discrepancy in preferring weight decay or dropout, our evaluation results consistently suggest that regularization is crucial to the imputation performance of autoencoders.

**Figure 9.**
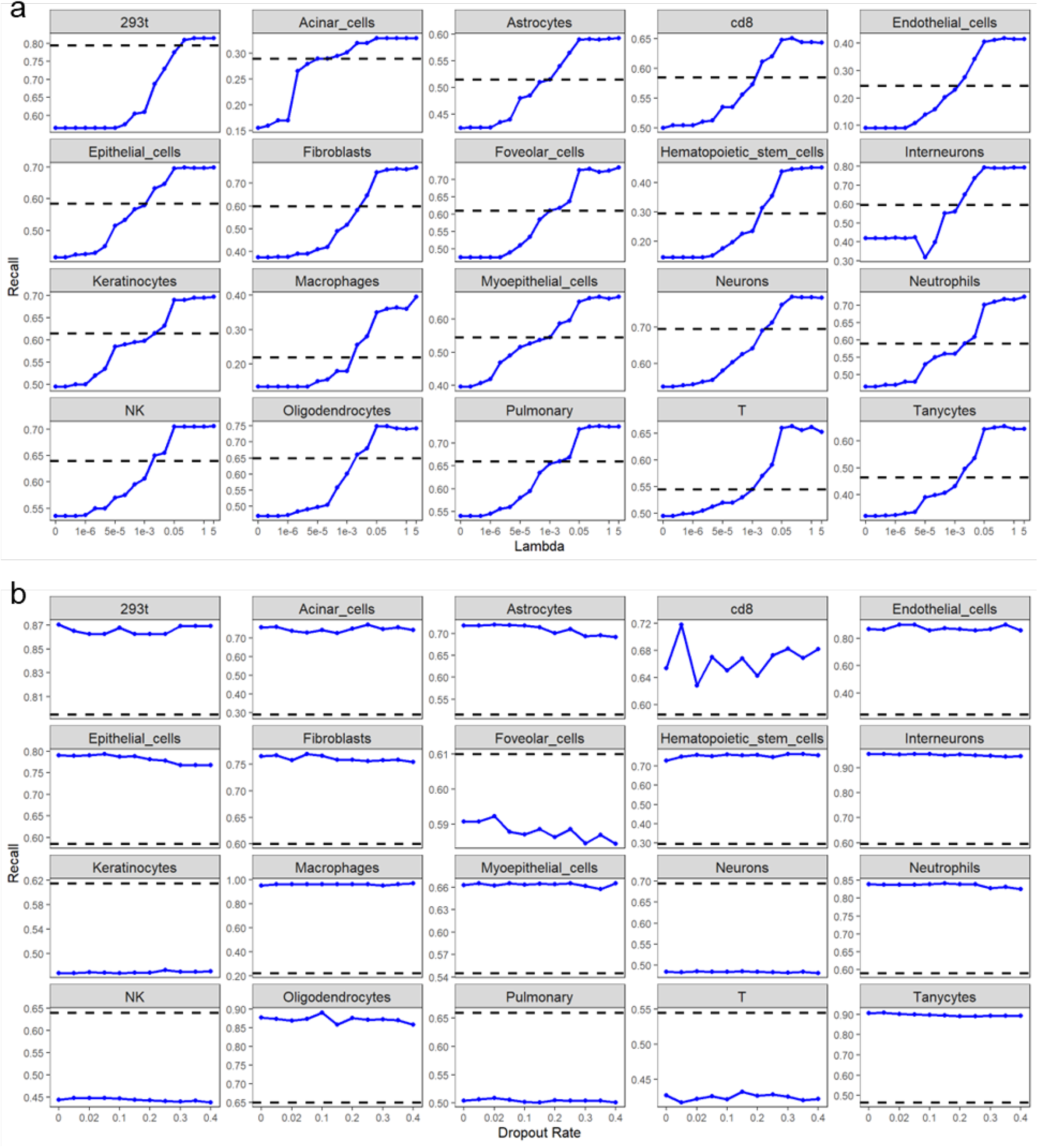
The impact of autoencoder regularization on DE gene identification accuracy measured by recall. **a**, Weight decay regularization. **b**, Dropout regularization. The dash lines show the recall without imputation. Each point is the average of the results obtained from five random seeds used for autoencoder training.

## Discussion

Sparsity is one of the major hurdles in scRNA-seq data analysis ^35^. To address the sparsity issue, more than 70 computational imputation methods have been developed, many of which are autoencoder-based ^36^, motivated by the success of neural networks in computer vision and natural language processing ^37^. Compared with traditional statistical and machine learning methods, autoencoder-based imputation methods have unique characteristics. First, autoencoder-based imputation methods require no assumptions on the underlying distribution of scRNA-seq data. This data-driven characteristic avoids the model specification bias in traditional methods. Second, autoencoder-based imputation methods can effectively handle large-scale scRNA-seq data by using innovative hardware, especially the GPU. Third, autoencoder-based imputation methods have high flexibility granted by their neural network design. With a carefully-designed loss function, they have incorporated multiple functionalities such as imputation, dimension reduction, and batch effect normalization ^38^.

However, how to optimize the design of autoencoders for scRNA-seq data remains a challenge. Successful applications of autoencoders in other fields, especially computer vision, rely on empirical studies that optimize the autoencoder design on massive datasets. Existing autoencoder-based imputation methods for scRNA-seq data follow certain guidelines (e.g., using the ReLU activation function) but ignore other guidelines (e.g., using a deep and narrow neural network) without comprehensive investigations. Our study fills in this gap, and our results confirm the deep learning field’s common wisdom that deeper and narrower autoencoders have better imputation performance. Meanwhile, we have an unexpected finding that sigmoid and tanh outperform ReLU and ReLU-modified forms as activation functions. This finding is consistent with another study, in which a neural network with the tanh activation function outperforms the ReLU counterpart in the cell-type classification task based on scRNA-seq data ^39^, reflecting the fact that scRNA-seq data are distinct from image data and thus need specific investigations.

Although our results favor deep and narrow neural networks, a previous study found that deep neural networks do not improve cell-type classification on scRNA-seq data ^40^. We hypothesize that this discrepancy is due to the two tasks’ different natures: our imputation task is a regression problem whose predictive targets are continuous gene expression values (after preprocessing), while the previous study’s task is a classification problem whose predictive targets are discrete cell-type labels. We hypothesize that, compared to classification, imputation requires deeper neural networks with a higher predictive capacity. Moreover, even though deeper autoencoders exhibit advantages in our study, the benefit of depth saturates when the number of layers surpasses 10, which is a much shallower architecture compared to the state-of-the-art deep neural networks with hundreds of layers used in computer vision. However, this discrepancy is consistent with the relationship between data complexity and model capacity. One image is typically a three-dimensional tensor (RGB channels × width × length) ^41^ and much more complex than one cell encoded in a one-dimensional vector in scRNA-seq data. Hence, it is unsurprising that scRNA-seq data need a shallower neural work than image data do.

Regarding the regularization strategy, we find that dropout excels in improving the overall imputation accuracy, and weight decay (using the *L*_2_ penalty) excels in improving the downstream cell clustering and DE gene analysis. Although dropout does not penalize all weight parameters, it randomly sets a certain proportion of weight parameters to zeros and thus can be interpreted as a stochastic *L*_1_ penalization. From this point of view, the *L*_1_ and *L*_2_ penalizations have complementary advantages in improving the overall imputation accuracy and the downstream cell clustering and DE gene analysis, respectively. We should also note that the three masking schemes are simplified approximations of, but are not themselves, the true missing mechanism, and the ultimate goal of imputation is to enhance downstream analyses. Therefore, weight decay may provide stronger benefits in real-world applications.

We observe that a similar autoencoder design exhibits distinctive imputation performance on scRNA-seq datasets of various characteristics. Datasets generated by 10X Genomics, e.g., jurkat, monocyte, and 293t, show higher overall imputation accuracy (Figure 2). Those datasets share some common characteristics, including high throughput and high zero rates (Supplementary Table S1). On the other hand, datasets generated by Smart-seq-total, Smart-seq2, and Fluidigm C1 show lower overall imputation accuracy (Figure 2). Those datasets have relatively low throughput with lower zero rates (Supplementary Table S1). When it comes to enhancing downstream cell clustering, datasets that contain a large number of ground-truth cell types, such as manno_human, chen, and lake, pose a greater challenge for autoencoders to enhance the separation of cell types. This is evident from Figure 6 and Supplementary Table S2. Even when the correct number of clusters is set in K-means clustering, it is more difficult for autoencoders to improve the cell clustering in those datasets. We also observe that autoencoders are less effective at improving DE gene identification on smaller datasets, such as Astrocytes, Macrophages, and Tanycytes (Supplementary Table S3). Compared with others, the sizes of those datasets typically range up to several hundred. Since the number of ground-truth DE genes is the same among all datasets, we suspect that the sample size plays a critical role in the effectiveness of autoencoder-based imputation methods.

Our findings indicate that the performance of autoencoder-based imputation methods is contingent upon the characteristics of the datasets and downstream tasks involved. While deeper autoencoders have the ability to capture non-linear and complex relationships in scRNA-seq data, their effectiveness may be limited in datasets with lower complexity. This observation is supported by evidence of performance saturation, as observed in Figures 2, 6, and 8, after reaching a certain number of layers. Considering the higher computational resources required for deeper autoencoders, it becomes crucial to balance the tradeoff between imputation performance and computational time during method design. Researchers must weigh the potential gains in imputation performance against the increased computational demands associated with deeper architectures.

Our study is conducted based on the original autoencoder. Some imputation methods use variants extended from the original autoencoder. For example, scVI is based on a variational autoencoder to learn a probabilistic latent space of the input data ^38^; scScope adopts an iterative autoencoder to impute data using many iterations ^42^; DCA uses the negative log-likelihood of a zero-inflated negative binomial model as the loss function to estimate the parameters of an autoencoder ^16^. Despite their differences, these autoencoder variants all have the encoder-decoder neural network structure and a non-linear activation function, with some including a regularization strategy. Therefore, our study can be easily generalizable to these autoencoder variants.

A major limitation of our study is that we adopt a greedy search strategy, instead of a global search strategy, to optimize the autoencoder design, due to the high computation complexity of autoencoder training. Hence, the autoencoder design we find as optimal in this study is not guaranteed to be globally optimal. One potential solution is to use advanced experimental design strategies, such as the space-filling design ^43,44^ and the fractional factorial design ^45,46^, to expand the search space so that the optimized design is closer to the global optimum. Another limitation is that we set all hidden layers to the same width (32, 64, 128, or 256) in each autoencoder, aiming to reduce the complexity of varying widths. Although there is no consensus on how the width heterogeneity across layers would affect an autoencoder’s learning capacity ^47^, it is possible that an autoencoder with hidden layers of different widths might have better imputation performance. We will leave these two improvements for a future study.

In summary, the performance of autoencoder-based imputation methods relies on key aspects of the autoencoder design, including the neural network architecture, activation function, and regularization strategy. Borrowing guidelines from practices in other fields cannot guarantee good performance on scRNA-seq data. Future methodological development should pay more attention to optimizing the autoencoder design and allow users to adjust the design for application needs.

## Methods

### Autoencoder for imputing scRNA-seq data

An autoencoder is a multi-layer neural network that aims to reconstruct the input data ^48^; it includes an encoder that embeds high-dimensional data to a low-dimensional latent space and a decoder that reconstructs the high-dimensional data from the low-dimensional embeddings (Figure 1a). Let *X* be a sparse scRNA-seq input matrix after appropriate preprocessing and normalization (see Methods section “Data preprocessing and normalization**”**). Let *Y* be the dense matrix output by an autoencoder. Both *X* and *Y* have *n* rows (cells) and *m* columns (genes). Denote *H*_*k*_, an *n*-by-*l*_*k*_ matrix, as the output of the *k*th hidden layer (with width *l*_*k*_) of the autoencoder, where *k* = 1, 2, …, ℎ, with ℎ as the total number of hidden layers. Then the output of the first hidden layer is

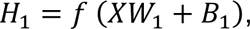

where *W*_1_ is an *m*-by-*l*_1_ weight matrix, and *B*_1_ is an *n*-by-*l*_1_ bias matrix with *n* identical rows. Here, *f* is an element-wise nonlinear activation function. Similarly, the output of the (*k* + 1)th hidden layer is

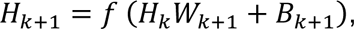

where *W*_*k*+1_ is an *l*_*k*_-by-*l*_*k*+1_ weight matrix, and *B*_*k*+1_ is an *n*-by-*l*_*k*+1_ bias matrix with *n* identical rows. Finally, the output of the autoencoder is

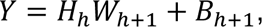

where *W*_ℎ+1_ is an *l*_ℎ_-by-*m* weight matrix and *B*_ℎ_ is an *n*-by-*m* bias matrix with *n* identical rows. Training the autoencoder means estimating (learning) the parameters in the weight matrices *W*_1_, *W*_2_, …, *W*_ℎ+1_ and the bias matrices *B*_1_, *B*_2_, …, *B*_ℎ+1_ by minimizing the mean squared error (MSE) between the input *X* and the output *Y* on the nonzero values of *X*. Let ***W*** be the set of weight matrices *W*_1_, *W*_2_, …, *W*_ℎ+1_ and ***B*** be the set of bias matrices *B*_1_, *B*_2_, …, *B*_ℎ+1_. Then the loss function is

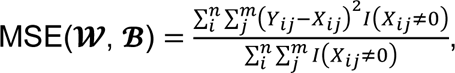

where *I*(·) is an indicator function. The sets of weight and bias matrices 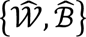 are estimated by

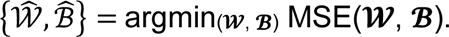

Since the minimization of MSE is a non-convex optimization problem ^47^, the backpropagation algorithm ^49^ is utilized to train the autoencoder (see Methods section “Autoencoder training and imputation”). In the imputation step, the zero entries in the input matrix *X* are replaced by their nonzero counterparts in the output matrix *Y*. That is, the imputed scRNA-seq data matrix 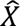 is calculated as

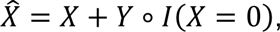

where ∘ is the element-wise product.

Several modifications to the original autoencoder have been made since the debut of autoencoder-based imputation methods. For example, DCA ^16^ models the scRNA-seq data by a negative binomial (NB) distribution with or without zero-inflation (ZI) (that is, ZINB or NB distribution) and learns the autoencoder by maximizing the likelihood of ZINB or NB calculated on the output *Y*; scVI ^38^ learns a variational autoencoder ^50^ by forcing each hidden unit in the hidden layer to follow a ZINB distribution; DeepImpute ^30^ learns an autoencoder by minimizing the weighted MSE, in which a gene’s weight is its expression level; LATE ^51^ considers cells or genes as observations, learns an autoencoder for each consideration, and selects the autoencoder with smaller MSE; scScope ^42^ learns an autoencoder by using the imputed data as input iteratively. Despite these modifications, all autoencoder-based imputation methods require the specification of the autoencoder design aspects we examine in this work.

### Regularization in autoencoder

Regularization is a technique for constraining the complexity of a machine learning model so that the model would have better generalizability ^52^. There are two regularization strategies, among others, commonly used to improve the imputation accuracy of autoencoders—weight decay ^47^ and dropout ^53^. Weight decay incorporates the *L*_2_ norm of weight parameters into the loss function to penalize large weights in an autoencoder. The weight and bias parameters under the weight decay 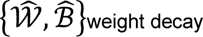 are given by

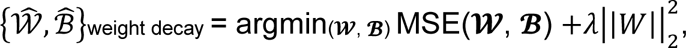

where 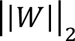 is the *L*_2_ norm of weight parameters, and *λ* is a tuning parameter that controls the degree of penalization.

Rather than penalizing the magnitudes of weight parameters, dropout regularization randomly sets a proportion of hidden units to zero in the training of an autoencoder. Dropout forces the autoencoder not to rely on particular hidden units and thus reduces overfitting ^53^. Specifically, suppose that *z*_*k*_ is a random vector with *l*_*k*_ elements, where *l*_*k*_ is the number of hidden units in the hidden layer *H*_*k*_. Each random variable in ***z_k_*** independently follows a Bernoulli distribution with a parameter *p*_*k*_∈ (0, 1). A matrix *Z*_*k*_ is constructed to have the same dimensions as those of the hidden layer matrix *H*_*k*_, and every row in *Z*_*k*_ is set to *z*_*k*_. Then in the training, the calculation of hidden layer *H*_*k*+1_ under dropout regularization is

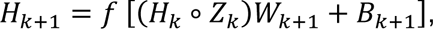

where ∘ is the element-wise product. Note that with a trained autoencoder, the calculation of hidden layers in the imputation step does not involve the dropout operation. In our analysis, we set *p*_1_ = *p*_2_ = ⋯ = *p*_ℎ_ = *p*, where ℎ is the total number of hidden layers. We call *p* the dropout rate in the following text.

### Three masking schemes for introducing artificial zeros

In scRNA-seq data, distinguishing biological zeros and non-biological zeros (i.e., missing values) is challenging without external information ^17,18^. Hence, to evaluate the overall imputation accuracy, we design three masking schemes to introduce artificial zeros to scRNA-seq data and measure the difference between the artificial zeros’ imputed values and original values (Figure 1b-d). The three masking schemes represent three typical assumptions of missing mechanisms in scRNA-seq data ^9^.

Random masking: we randomly mask (set values to zero) 50% nonzero entries in the scRNA-seq data matrix. Random masking means that the missing mechanism is independent of the gene expression levels, and it has been widely used in previous work to evaluate the imputation accuracy ^18^.

Median masking: we mask the nonzero entries less than or equal to the overall median in the scRNA-seq data matrix. Median masking assumes complete dependence of the missing mechanism on the gene expression levels.

Double exponential masking: we assume that a gene’s probability of having missing values depends on the gene’s mean expression level across all cells. Hence, lowly expressed genes are more likely to have missing values than highly expressed genes. Specifically, for gene *j*, let 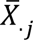 be the mean expression level (natural-log-plus-one-transformed read count) of nonzero values across all cells and *p*_*j*_ be the probability of missing values. Then gene *j*’s probability of having missing values is modeled by a double exponential function ^9^

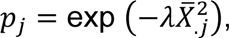

where *λ* is a parameter estimated from scRNA-seq data across all genes, using *p*_*j*_’s calculated as gene *j*’s proportion of zero counts and 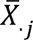 as gene *j*’s average of nonzero expression levels. Let *Z*_*ij*_ be a random variable that indicates whether to mask the nonzero expression of gene *j* in cell *i*. Then *Z*_*ij*_ ∼ *Bernoulli* (*p_i_*). If *Z*_*ij*_ = 1, then the nonzero expression of gene *j* in cell *i* will be masked. The value of *λ* is determined such that 50% nonzero entries in the scRNA-seq data matrix are masked.

A previous study showed that artificial zeros introduced by the three masking schemes exhibited different impacts on scRNA-seq data analysis ^9^. Although random masking has been widely used to evaluate imputation accuracy, its assumption is unrealistic. In contrast, median masking is an extreme way of modeling the observation that lowly expressed genes have more zeros than highly expressed genes have ^15,54^. Between the two masking schemes, double exponential masking uses a probabilistic model to reflect this observation at the individual gene level. Together, the three masking schemes provide a comprehensive evaluation of imputation accuracy from different perspectives.

### Data preprocessing and normalization

All real and synthetic scRNA-seq datasets used in this study are cell-by-gene count matrices. They are preprocessed and normalized in three steps. First, we remove the genes expressed in fewer than three cells and the cells having fewer than 200 genes expressed. Second, we divide each count by its cell library size (i.e., the cell’s total count) and then multiply it by 10000 (library size normalization). Then we add one to the normalized count and apply the natural-log transformation. Third, we select 2000 highly variable genes using the `vst` method implemented in the `FindVariableFeatures` function of the `Seurat` package ^33^ (v4.0). After preprocessing and normalization, the dimensions of all scRNA-seq data matrices are cell number × 2000. Note that the preprocessing and normalization steps apply to the pre-imputed datasets only. The imputed datasets, when used as the input of cell clustering and DE gene analysis, will not go through the preprocessing and normalization steps.

### Training of autoencoders and imputation

The training of autoencoders is implemented using the `Pytorch` deep learning library ^55^ (v 1.8.1) on a server with two Intel Xeon E5-2687W v4 CPUs, 256GB memory, an Nvidia GeForce RTX 2080 Ti GPU, and Ubuntu 18.04 system. After preprocessing, normalization, and masking (masking is only necessary for the evaluation of overall imputation accuracy), we split each dataset’s cells into a training set (80% of cells) and a validation set (20% of cells). We utilize the Adam optimization algorithm ^21^ to train the autoencoder with a 0.001 learning rate and a 64 batch size. After every epoch of training on the training set, we impute the validation set using the current autoencoder and calculate the MSE between the original nonzero values in the validation set and their corresponding imputed values. We continue the training until either the MSE does not decrease over 20 epochs or the total number of epochs surpasses 10000. In the imputation step, the trained autoencoder accepts the preprocessed and normalized scRNA-seq data matrix as input (with the dimension as cell number × 2000) and outputs a data matrix of the same dimension. The final imputed data matrix is constructed by replacing the zero values in the input matrix with their counterparts in the output matrix. The nonzero values in the input matrix remain in the final imputed data matrix.

### Calculation of imputation normalized root mean squared error (NRMSE)

Denote by *X* the sparse scRNA-seq input matrix after preprocessing and normalization; 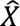 is the imputed scRNA-seq data matrix; *M* is the set of masked entries in the scRNA-seq data matrix. The MSE between the imputed and original values of the masked entries, MSE_mask_, is calculated as

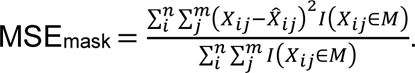

Denote the mean masked values 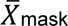 as

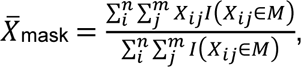

then the imputation NRMSE is calculated as

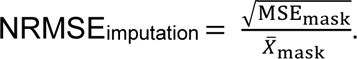

### Activation functions

We evaluate seven activation functions in this study, including logistic function (sigmoid), hyperbolic tangent function (tanh), rectified linear unit (ReLU), leaky ReLU (with two hyperparameters), exponential linear units (ELU), and scaled exponential linear units (SELU). An activation function accepts a linear combination of the outputs from the last layer as input and applies a nonlinear transformation to the linear combination. All activation functions take a real-valued scalar as input and output a real-valued scalar.

The sigmoid activation function is a bounded differentiable function with positive and continuous derivatives. Its value ranges from 0 to 1. The function form of sigmoid is

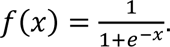

The tanh activation function is also a bounded differentiable function with positive and continuous derivatives. Its value ranges from −1 to 1. The functional form of tanh is

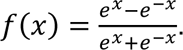

The ReLU activation function conducts a threshold operation that outputs zero for a negative input and acts as an identity function for a positive input. Its value ranges from 0 to +∞, and it has discrete derivatives. The functional form of ReLU is

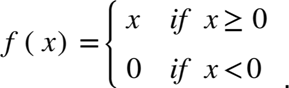

The leaky ReLU activation function modifies ReLU by introducing a small negative slope when the input is negative. Its value ranges from −∞ to +∞, and it has discrete derivatives. The functional form of leaky ReLU is

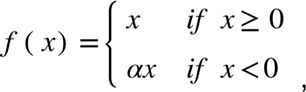

where *α* > 0 is a hyperparameter. In our analysis, we set *α* to 0.01 and 0.2 — the default values in two popular deep learning libraries `Pytorch` ^55^ and `TensorFlow` ^56^.

The ELU activation function replaces the linear negative part of leaky ReLU with an exponential function. Its value ranges from −∞ to +∞, and it has discrete derivatives. The functional form of ELU is

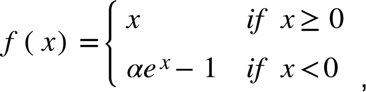

where *α* > 0 is a hyperparameter. In our analysis, we set *α* to 1 — the default value in `Pytorch`.

The SELU activation function further adds a scale factor to ELU and changes the constant in the negative part. Its value ranges from −∞ to +∞, and it has discrete derivatives. The functional form of ELU is

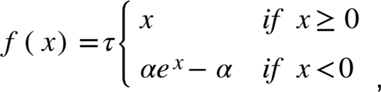

where *τ* > 0 and *α* > 0 are predefined parameters with *τ* = 1.05 and *α* = 1.67.

### Weight decay and dropout parameter settings

When evaluating the impact of weight decay regularization on the imputation accuracy, we set the hyperparameter *λ* to nine values, including 0, 1e-7, 5e-7, 1e-6, 5e-6, 1e-5, 5e-5, 1e-4, and 5e-4. Larger *λ* values indicate stronger regularization imposed on the autoencoder, and *λ* = 0 means no regularization. When evaluating the impact of weight decay regularization on the cell clustering and DE gene analysis, we set the hyperparameter *λ* to 17 values, including 0, 1e-7, 5e-7, 1e-6, 5e-6, 1e-5, 5e-5, 1e-4, 5e-4, 5e-3, 0.01, 0.05, 0.1, 0.5, 1, and 5. When evaluating the impact of dropout regularization on the imputation accuracy, cell clustering, and DE gene analysis, we set the dropout rate *p* to 11 values, including 0, 0.01, 0.02, 0.05, 0.1, 0.15, 0.2, 0.25, 0.3, 0.35, and 0.4. Larger *p* values indicate stronger regularization imposed on the autoencoder, and *p* = 0 means no regularization.

### Cell clustering analysis

We use the function `kmeans` in R programming language to conduct K-means clustering on the pre-imputed and imputed scRNA-seq datasets (Supplementary Table S2). We set the parameter `centers` (the number of clusters *k* in the K-means clustering) to the correct number of cell types in each dataset. We set the parameter `nstart` to 25, which repeats the clustering 25 times by randomly selecting 25 sets of initial cluster centers and returns the result with the minimum sum of pairwise distances within clusters ^57^. The dimension of input data matrices for K-means clustering is cell number × 2000 without further dimension reduction (that is, each cell is a 2000-dimensional vector). Note that before clustering, pre-imputed datasets are preprocessed by following the procedure described in the section “Data preprocessing and normalization.” The imputed datasets are directly clustered by K-means clustering.

We use adjusted rand index (ARI) and adjusted mutual information (AMI) to measure the performance of cell clustering. Let *U* = {*u*_1_, *u*_2_, …, *u*_*c*_} be the true partition of *c* classes and *V* = {*v*_1_, *v*_2_, …, *v*_*c*_} be the partition obtained by K-means clustering. Let *n*_*i*_ and *m*_*j*_ be the numbers of observations in class *u*_*i*_ and cluster *v*_*j*_, respectively. Let *r*_*ij*_ be the number of observations in both class *u*_*i*_ and cluster *v*_*j*_. The ARI is calculated as

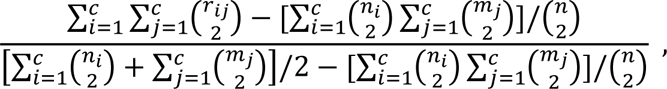

where *n* is the total number of observations and 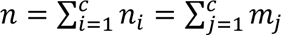. The AMI is calculated as

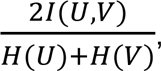

where *I*(*U*, *V*) is the mutual information of *U* and *V*, and *H*(*U*) and *H*(*V*) are the entropies of *U* and *V* respectively ^58^. We utilize the functions `ARI` and `AMI` in R package `aricode` to calculate ARI and AMI, respectively.

### Simulation of synthetic scRNA-seq data

We utilize simulator scDesign ^19^ to generate 20 synthetic scRNA-seq data with ground-truth DE genes. The 20 real datasets for training scDesign (Supplementary Table S3) are preprocessed by following the procedure described in the section “Data preprocessing and normalization.” For each real dataset, we execute function `design_data` in R package `scDesign` to simulate one synthetic dataset. Each synthetic dataset contains two cell types with 1000 cells per type, and 10% genes are differentially expressed between the two cell types. The sequencing depth of each synthetic dataset is equal to the median cell library size of the corresponding real dataset multiplied by the cell number (2000). Other parameters of function `design_data` are set to their default values. All synthetic datasets are count matrices with dimensions as cell number × 2000.

### DE gene analysis

We conduct DE gene analysis on the aforementioned 20 synthetic datasets and their imputed counterparts. For pre-imputed synthetic datasets, the gene expression counts of each cell are divided by the total counts of that cell (library size) and then multiplied by 10000 (library size normalization). The results are then added by one before being natural-log transformed. We utilize the function `FindMarkers` in R package `Seurat` to identify the DE genes between the two cell types. We set the parameter `test.use` to `“MAST”` and identify genes with Bonferroni-corrected p-values under 0.05 as DE genes. Based on the ground-truth DE genes, we calculate the precision, recall, and TNR for each pre-imputed synthetic dataset and the corresponding imputed dataset.

### Sensitivity analysis with varying numbers of highly variable genes

We performed a sensitivity analysis by varying the number of highly variable genes to assess the robustness of our major findings. Specifically, we examined the impact of autoencoder architecture, activation function, and regularization on imputation accuracy using random masking. Two scenarios were considered, where the highly variable genes were set to 1000 and 3000, respectively. The results of this analysis are summarized in Supplementary Figures S12–S19. Notably, our investigation revealed that our major findings remained consistent across different numbers of highly variable genes. This suggests that the observed trends and conclusions are not dependent on the specific number of highly variable genes considered in the analysis.

## Data and Code Availability

The source code implements this study is available at GitHub repository https://github.com/xnnba1984/Benchmarking-the-Autoencoder-Design-for-Imputing-Single-Cell-RNA-Sequencing-Data.

The datasets used in this study can be found at Zenodo repository https://zenodo.org/record/7504311#.Y7ei0naZMuK

## Supplementary Figures and Tables

**Supplementary figure S1.**
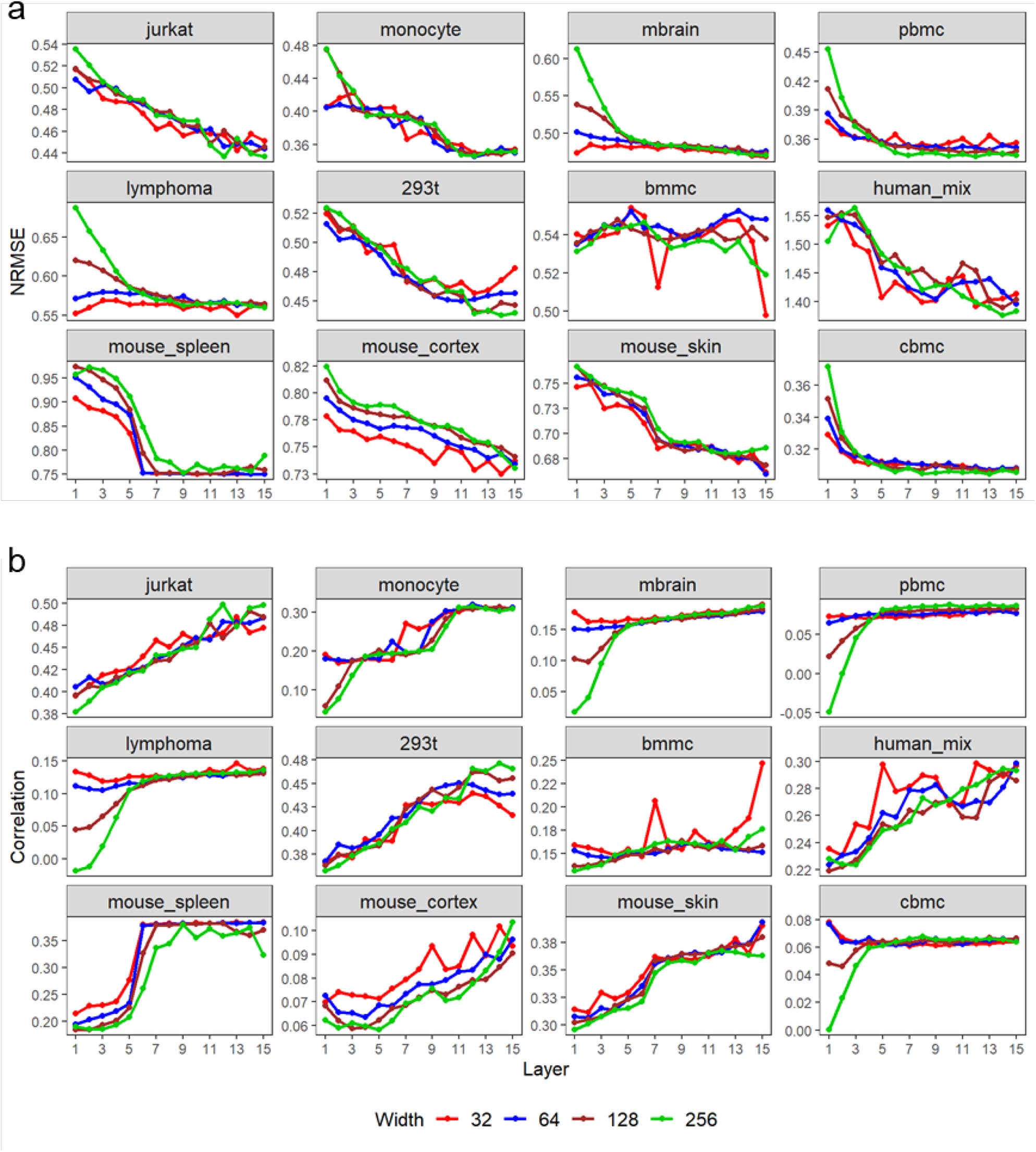
The impact of neural network depth and width on the imputation NRMSE (**a**) and the imputation correlation (**b**) based on the double exponential masking scheme. Each point is the average of the results obtained from 10 random seeds used for autoencoder training.

**Supplementary figure S2.**
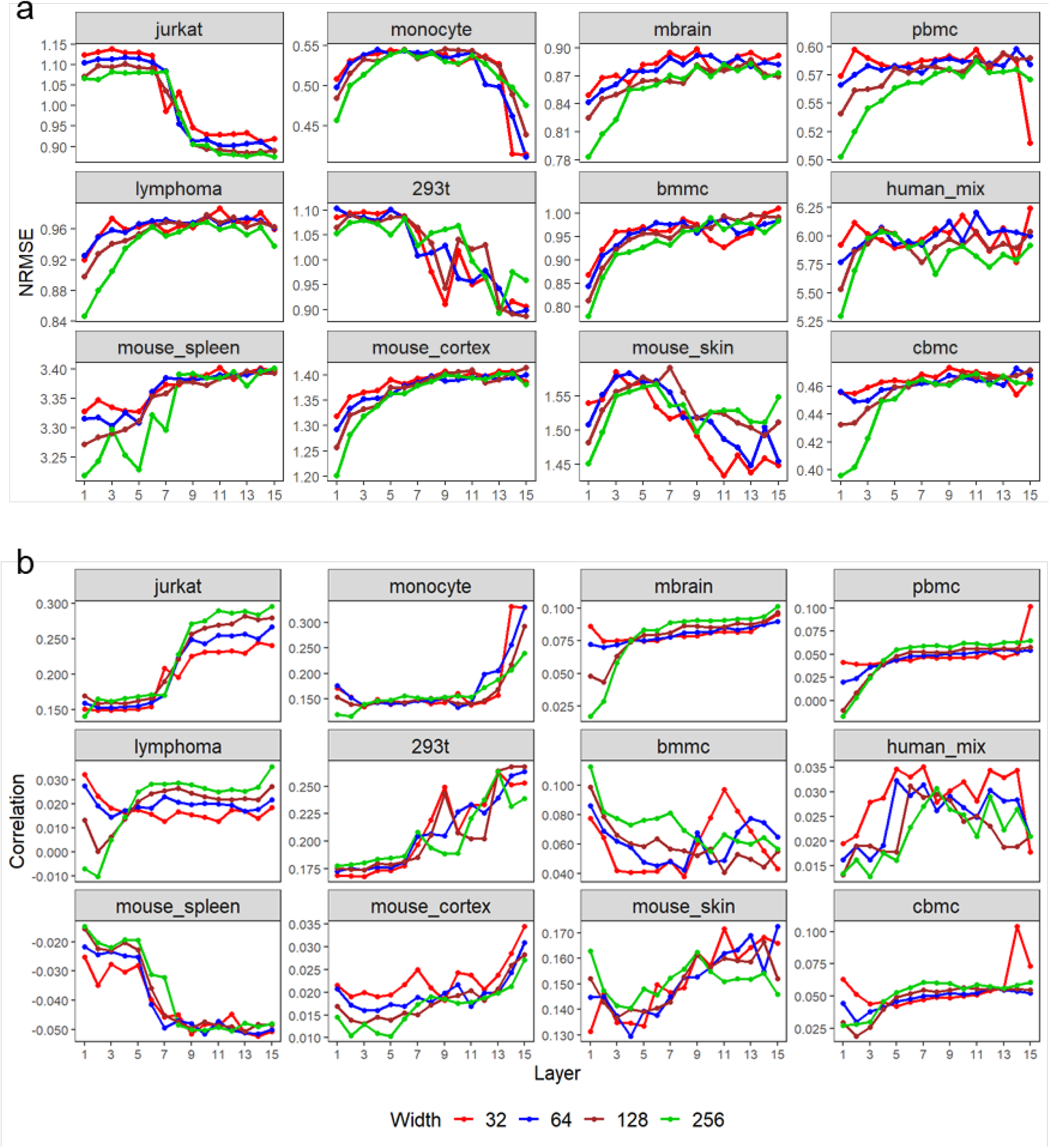
The impact of neural network depth and width on the imputation NRMSE (**a**) and the imputation correlation (**b**) based on the median masking scheme. Each point is the average of the results obtained from 10 random seeds used for autoencoder training.

**Supplementary Figure S3.**
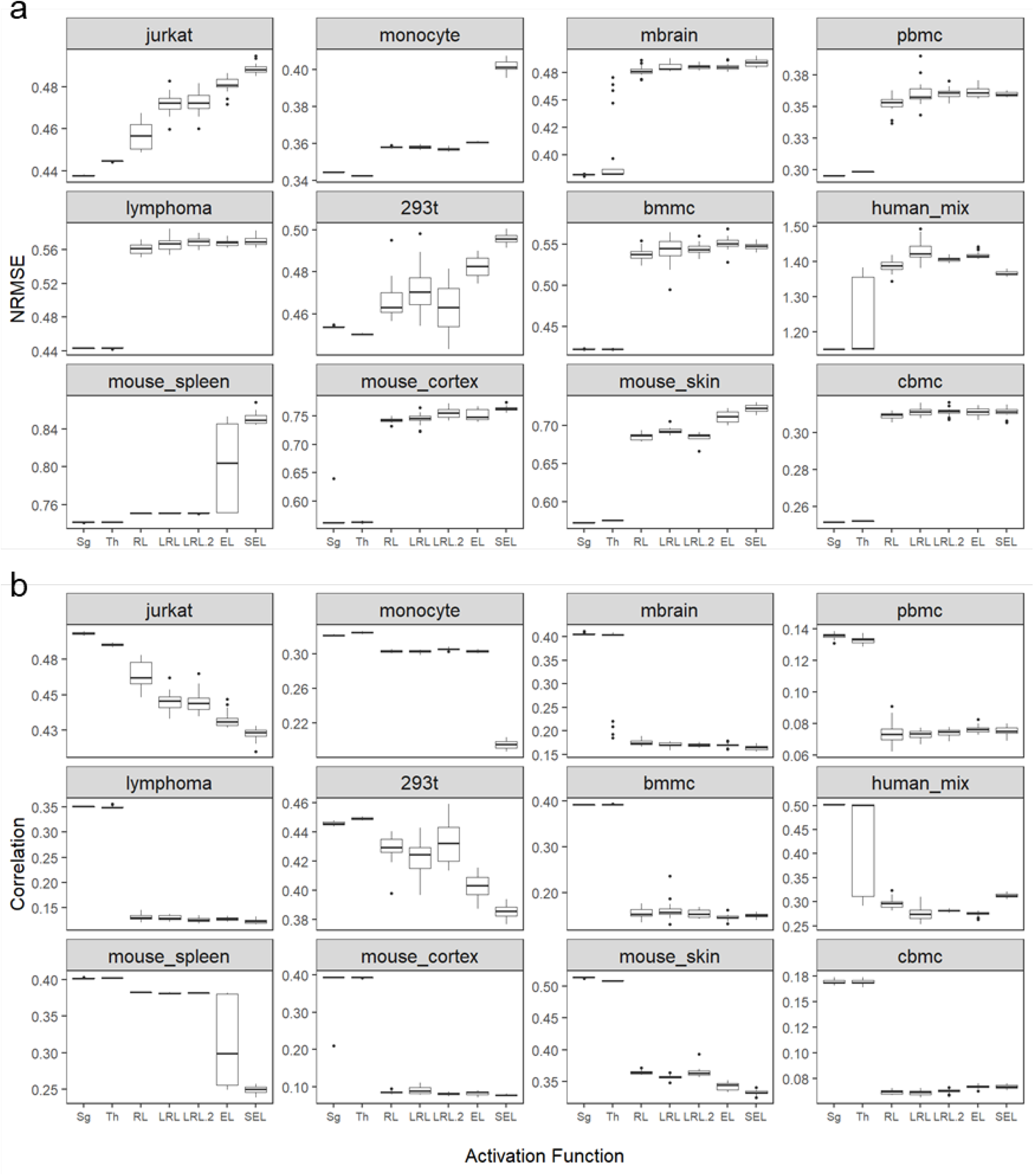
The impact of activation functions on the imputation NRMSE (**a**) and the imputation correlation (**b**) based on the double exponential masking scheme. Sg: sigmoid; Th: tanh; RL: ReLU; LRL: LeakyReLU (*α* = 0.01); LRL.2: LeakyReLU (*α* = 0.2); EL: ELU; SEL: SELU. Each boxplot shows the results obtained from 20 random seeds used for autoencoder training.

**Supplementary Figure S4.**
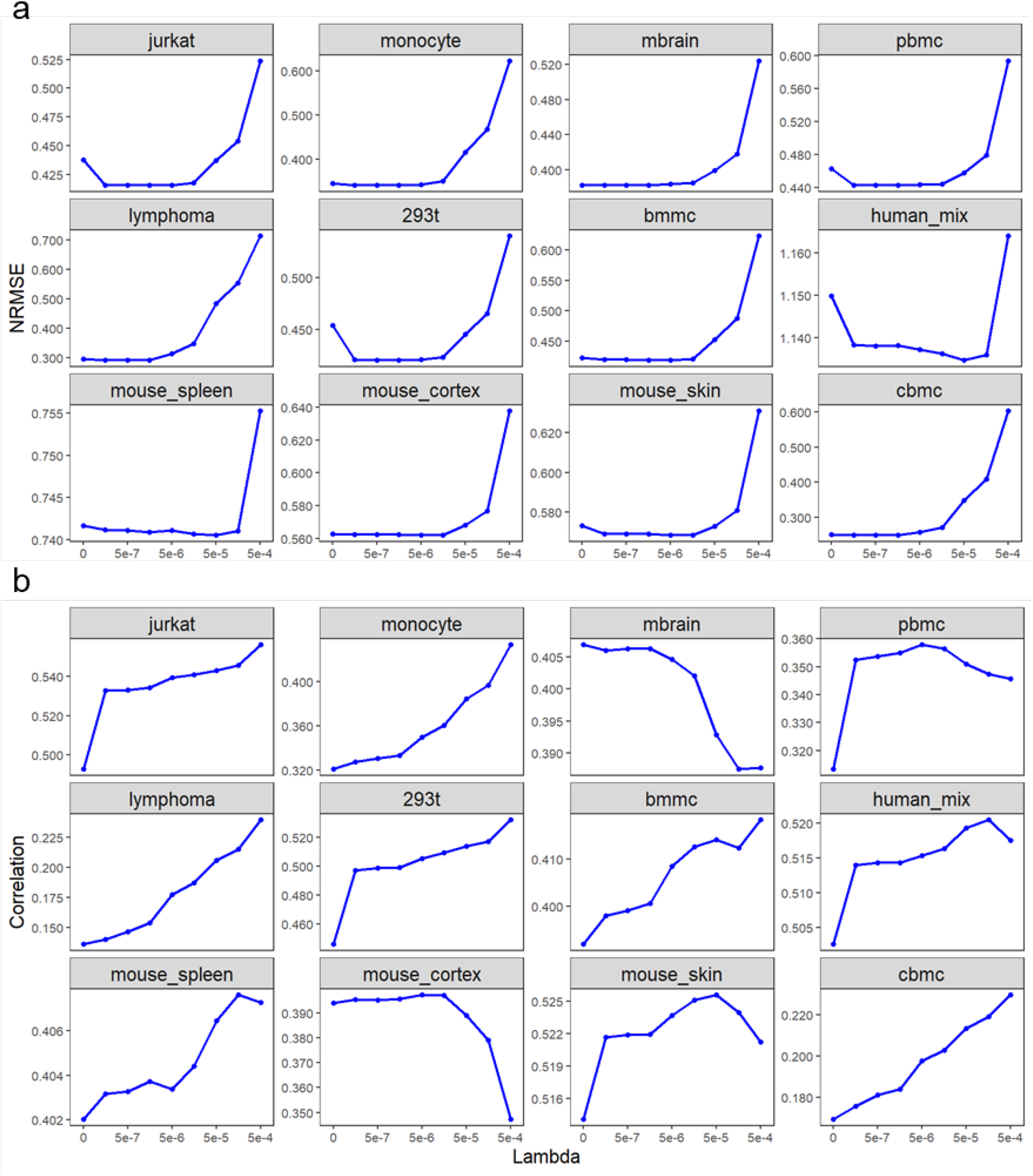
The impact of weight decay regularization on the imputation NRMSE (**a**) and the imputation correlation (**b**) based on the double exponential masking scheme. Each point is the average of the results obtained from 10 random seeds used for autoencoder training.

**Supplementary Figure S5.**
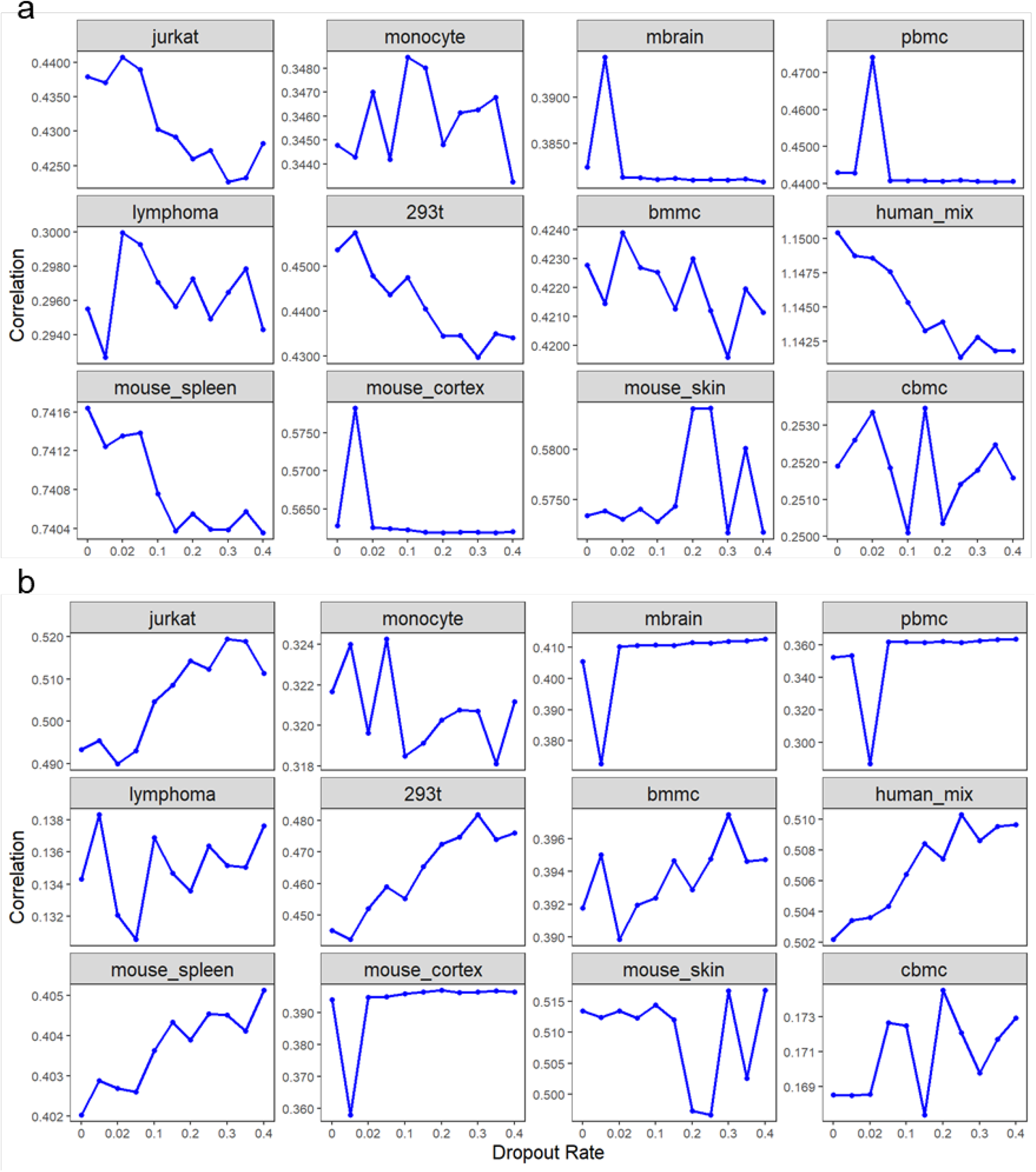
The impact of dropout regularization on the imputation NRMSE (**a**) and the imputation correlation (**b**) based on the double exponential masking scheme. Each point is the average of the results obtained from 10 random seeds used for autoencoder training.

**Supplementary Figure S6.**
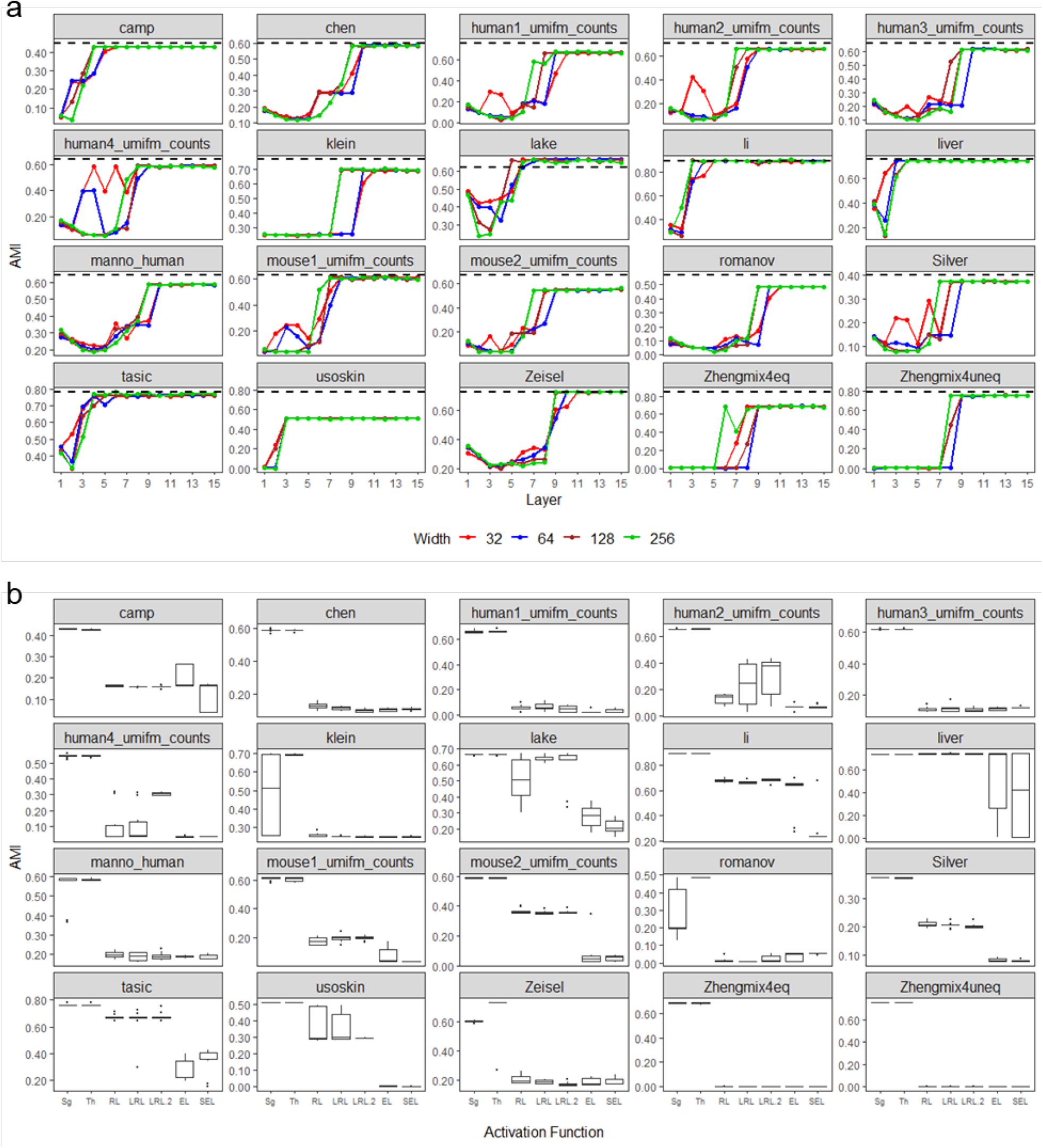
The impact of autoencoder design on cell clustering accuracy measured by AMI. **a,** Depth and width. **b**, Activation function. The dash lines in (**a**) show the cell clustering performance without imputation. Each point in (**a**) is the average of the results obtained from five random seeds used for autoencoder training. Each boxplot in (**b**) shows the results obtained from 10 random seeds used for autoencoder training.

**Supplementary Figure S7.**
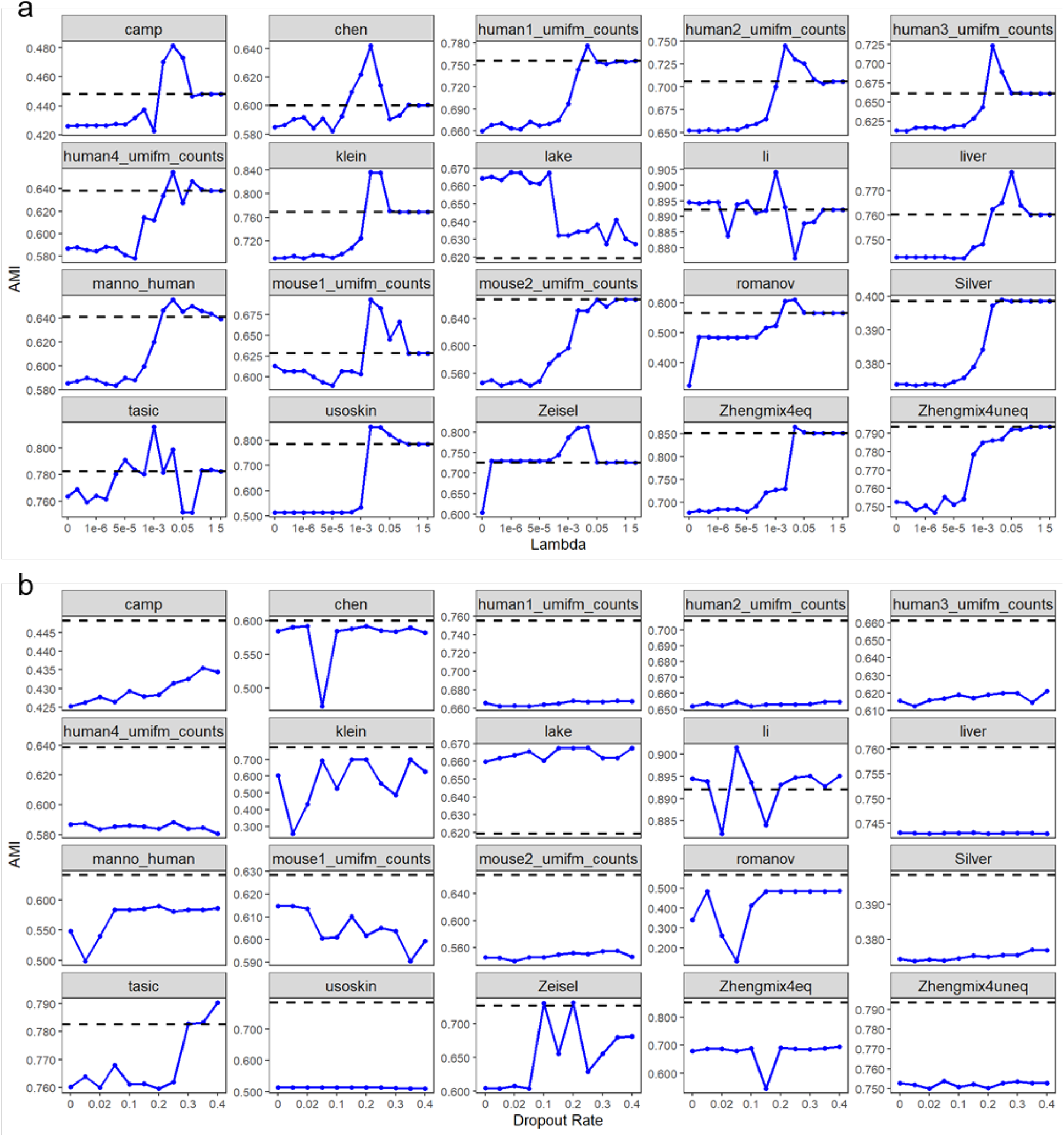
The impact of autoencoder regularization on cell clustering accuracy measured by AMI. **a**, Weight decay regularization. **b**, Dropout regularization. The dash lines show the cell clustering performance without imputation. Each point is the average of the results obtained from five random seeds used for autoencoder training.

**Supplementary Figure S8.**
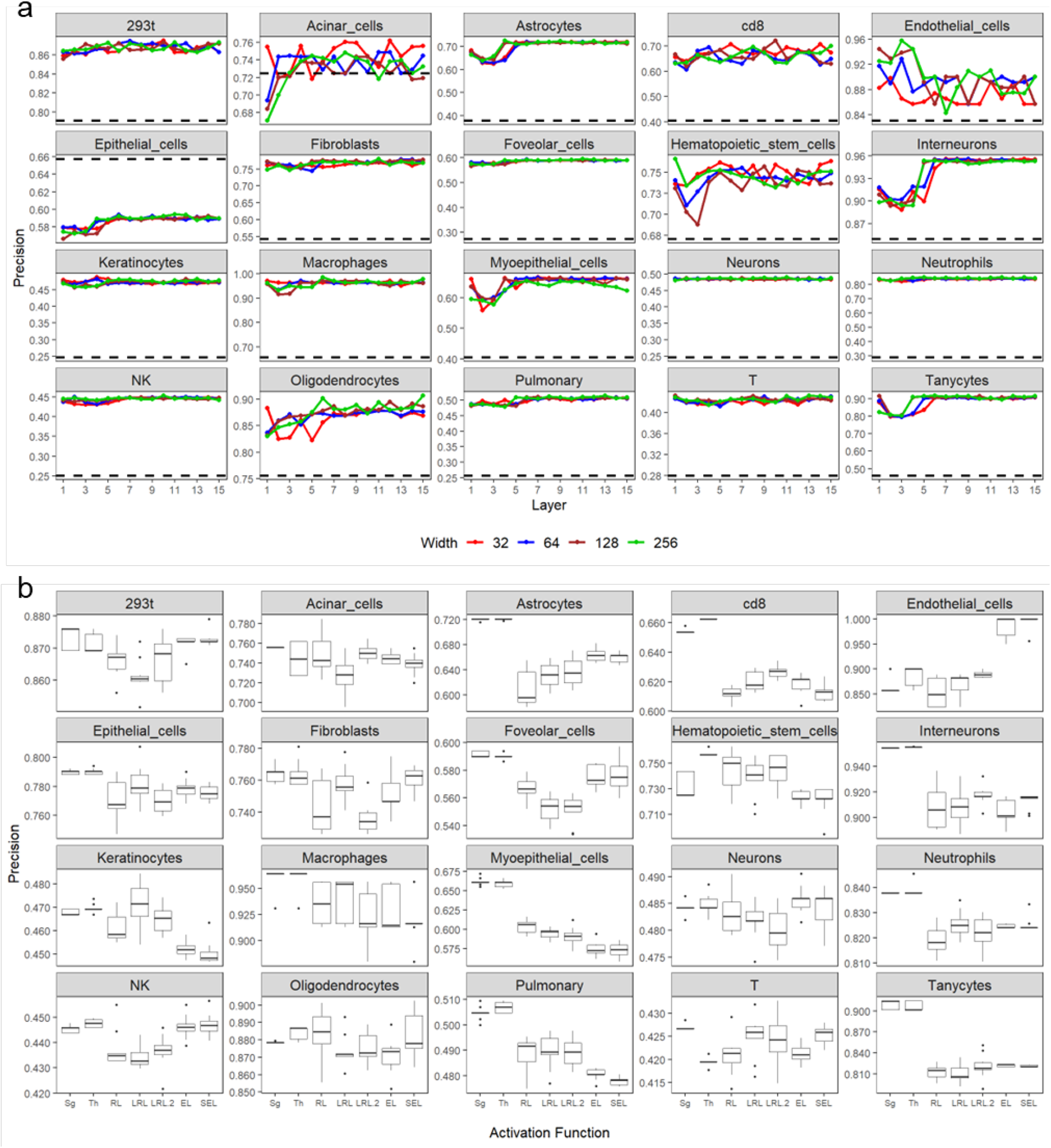
The impact of autoencoder design on DE gene analysis measured by precision. **a,** Depth and width. **b**, Activation function. The dash lines in (**a**) show the precision without imputation. Each point in (**a**) is the average of the results obtained from five random seeds used for autoencoder training. Each boxplot in (**b**) shows the results obtained from 10 random seeds used for autoencoder training.

**Supplementary Figure S9.**
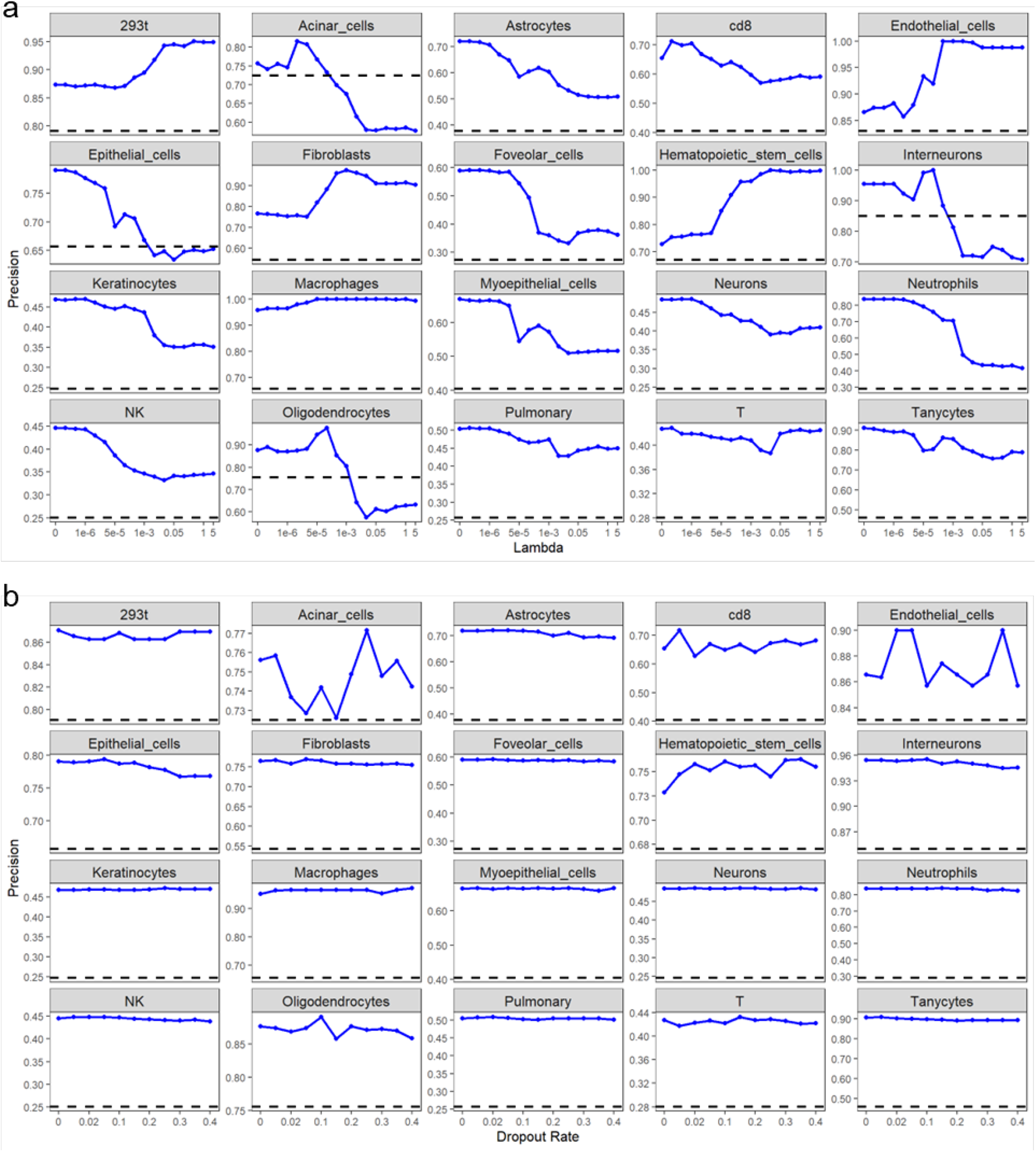
The impact of autoencoder regularization on DE gene analysis measured by precision. **a**, Weight decay regularization. **b**, Dropout regularization. The dash lines show the precision without imputation. Each point is the average of the results obtained from five random seeds used for autoencoder training.

**Supplementary Figure S10.**
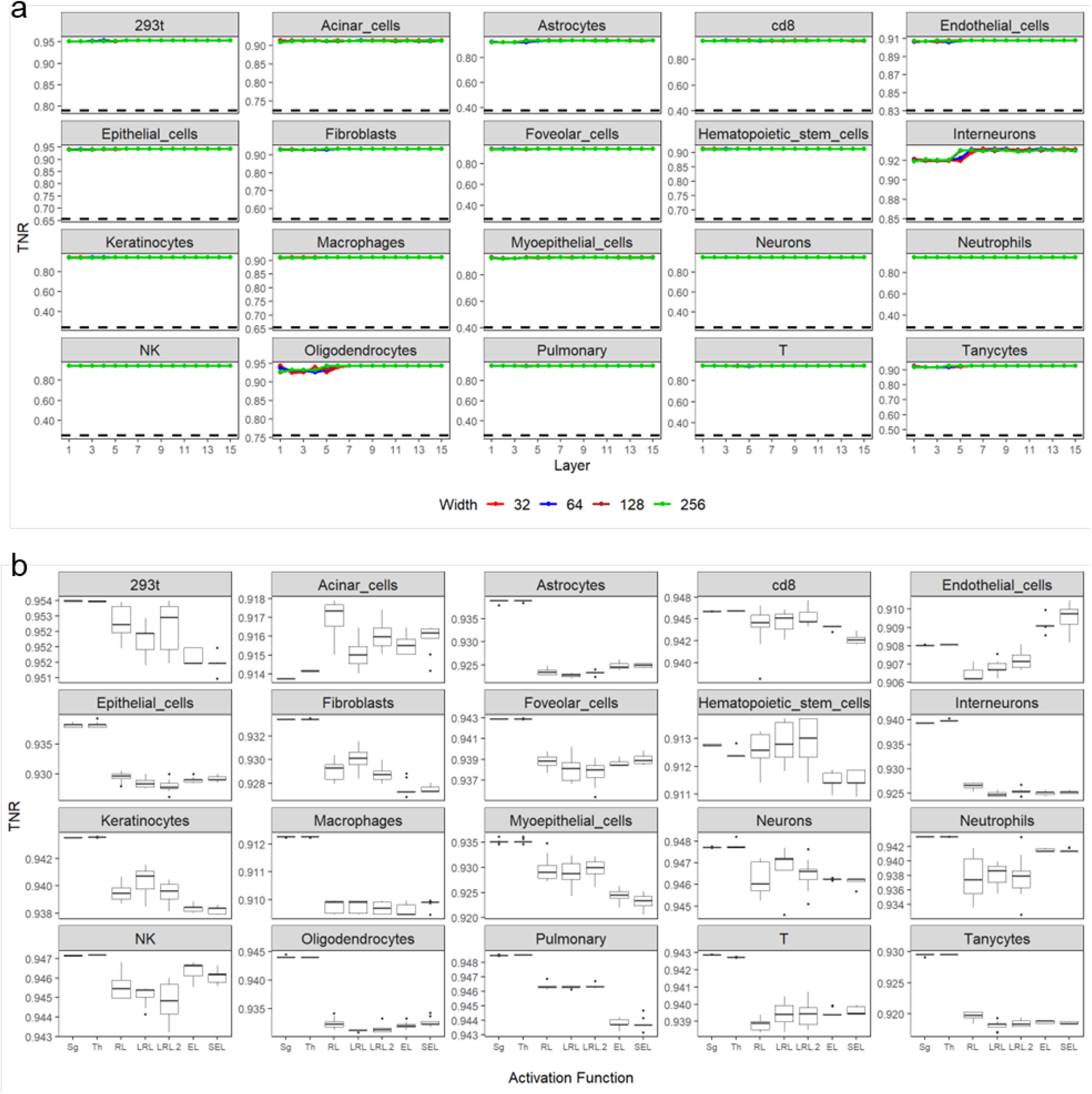
The impact of autoencoder design on DE gene analysis measured by TNR. **a,** Depth and width. **b**, Activation function. The dash lines in (**a**) show the TNR without imputation. Each point in (**a**) is the average of the results obtained from five random seeds used for autoencoder training. Each boxplot in (**b**) shows the results obtained from 10 random seeds used for autoencoder training.

**Supplementary Figure S11.**
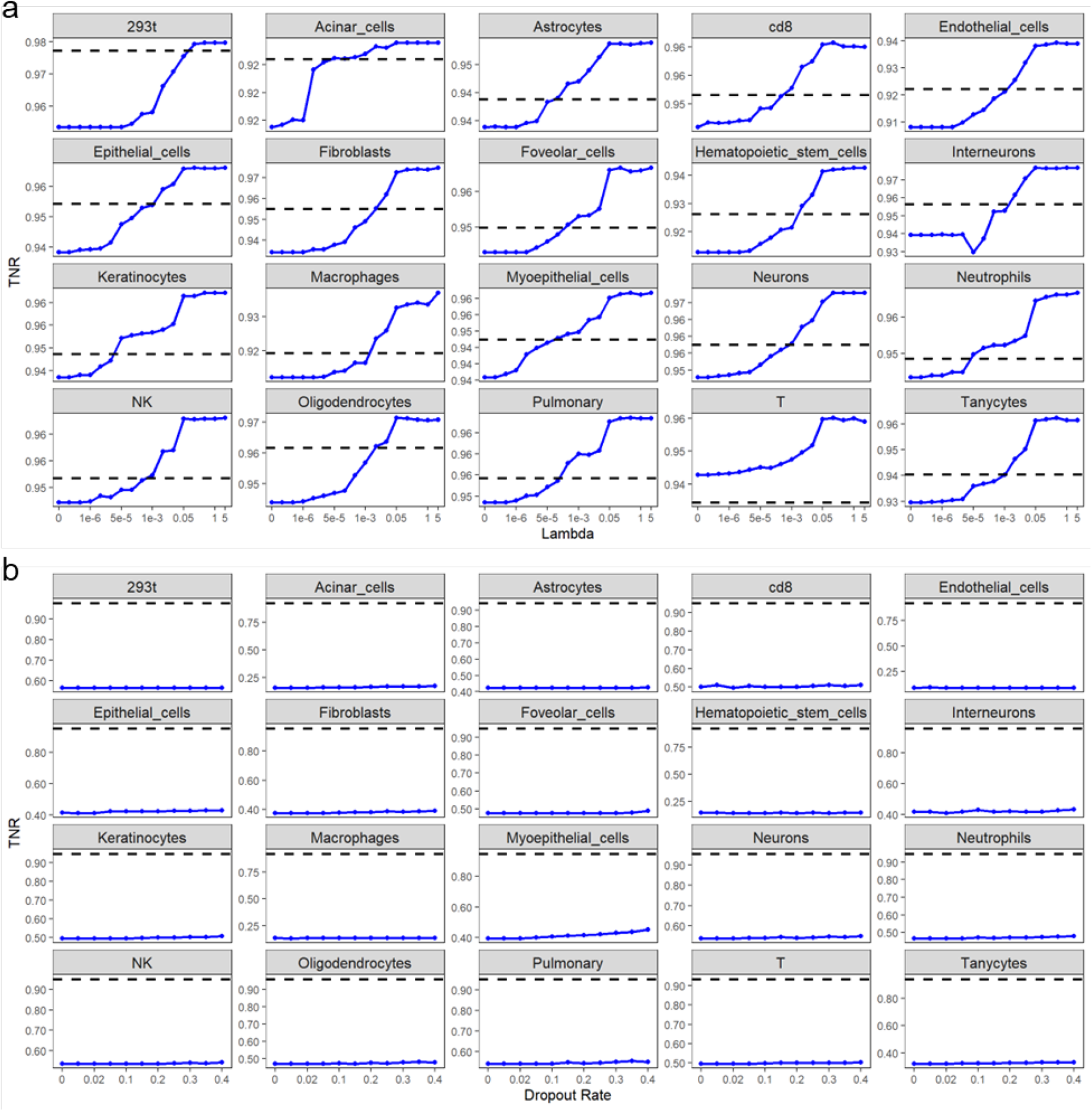
The impact of autoencoder regularization on DE gene analysis measured by TNR. **a**, Weight decay regularization. **b**, Dropout regularization. The dash lines show the TNR without imputation. Each point is the average of the results obtained from five random seeds used for autoencoder training.

**Supplementary Figure S12.**
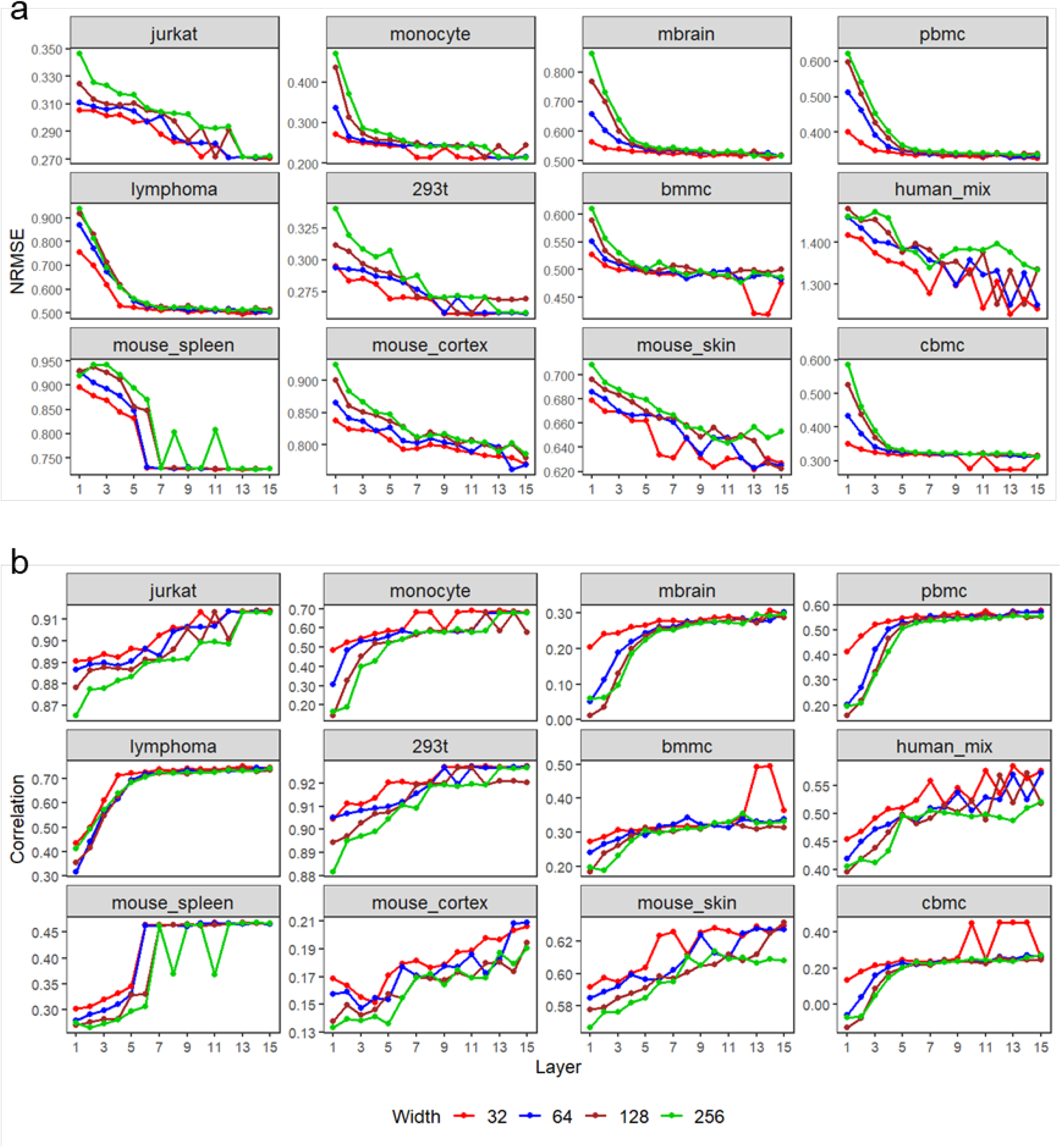
The impacts of autoencoder depth and width on the imputation NRMSE (**a**) and the imputation correlation (**b**) based on the random masking scheme. The number of highly variable genes is set to 1000.

**Supplementary Figure S13.**
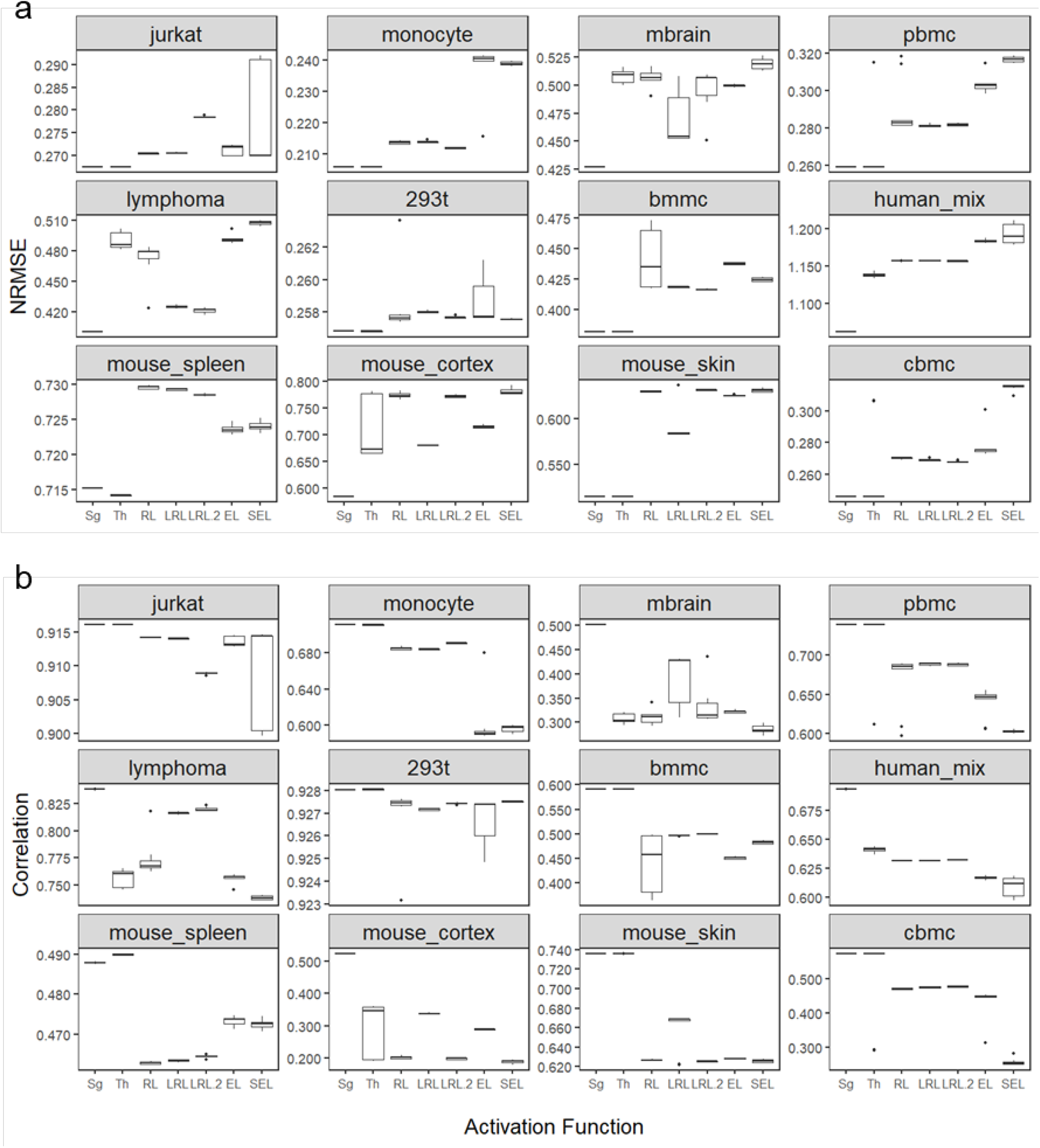
The impact of activation functions on the imputation NRMSE (**a**) and the imputation correlation (**b**) based on the random masking scheme. Sg: sigmoid; Th: tanh; RL: ReLU; LRL: LeakyReLU (=0.01); LRL.2: LeakyReLU (=0.2); EL: ELU; SEL: SELU. The number of highly variable genes is set to 1000.

**Supplementary Figure S14.**
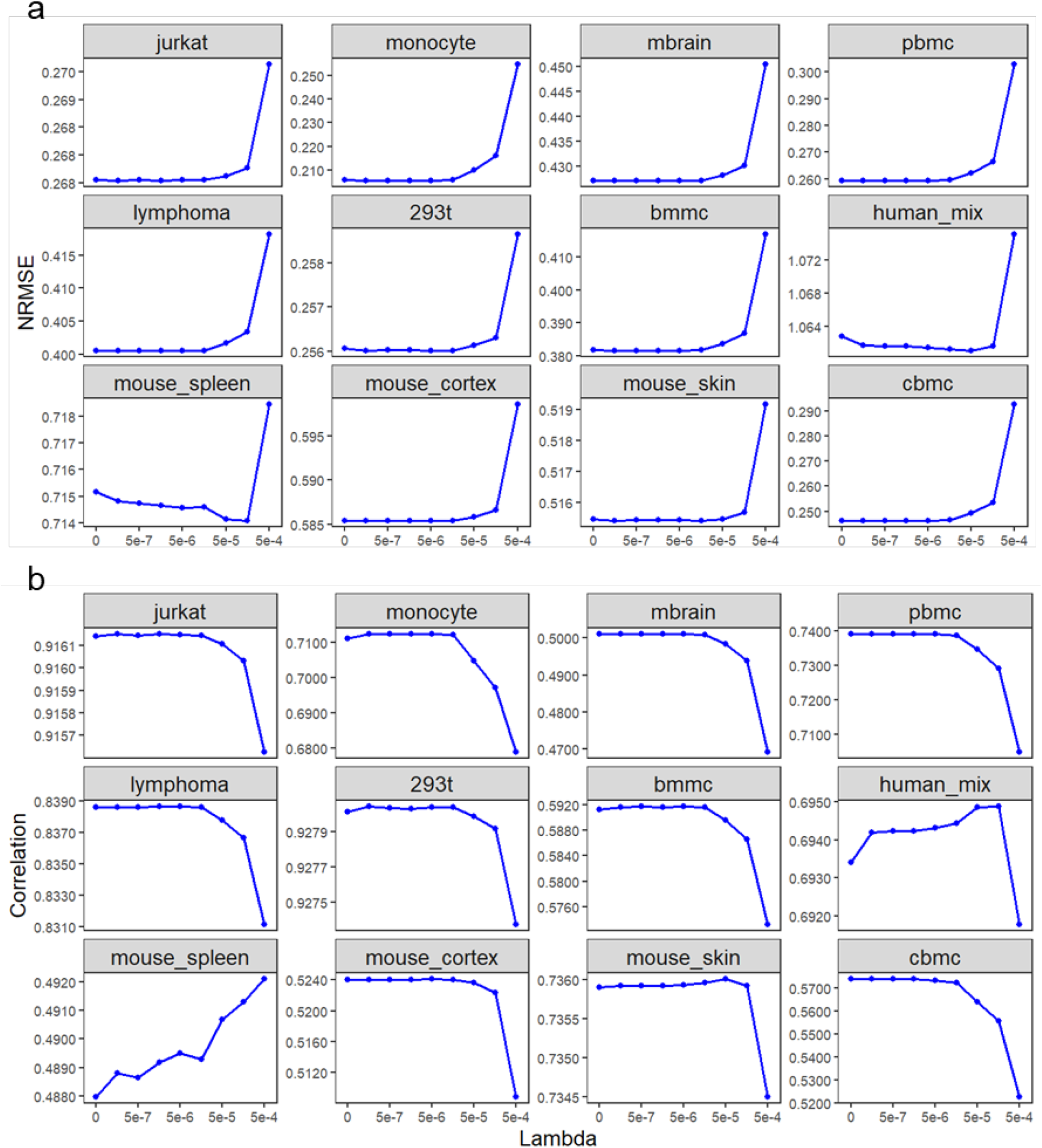
The impact of weight decay regularization on the imputation NRMSE (**a**) and the imputation correlation (**b**) based on the random masking scheme. The number of highly variable genes is set to 1000.

**Supplementary Figure S15.**
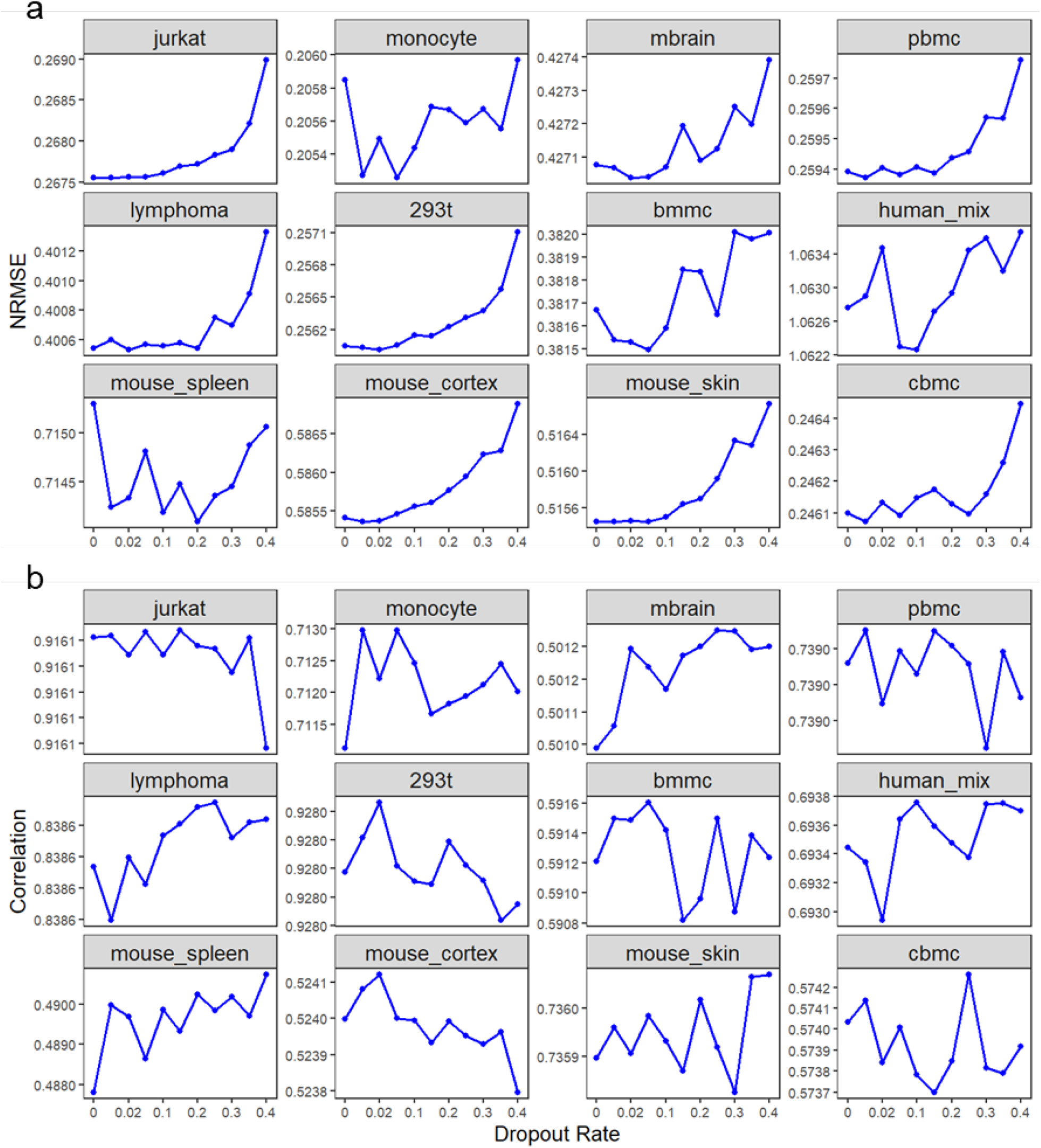
The impact of dropout regularization on the imputation NRMSE (**a**) and the imputation correlation (**b**) based on the random masking scheme. The number of highly variable genes is set to 1000. Some labels on the y-axis are identical due to the low accuracy variability under different dropout rates.

**Supplementary Figure S16.**
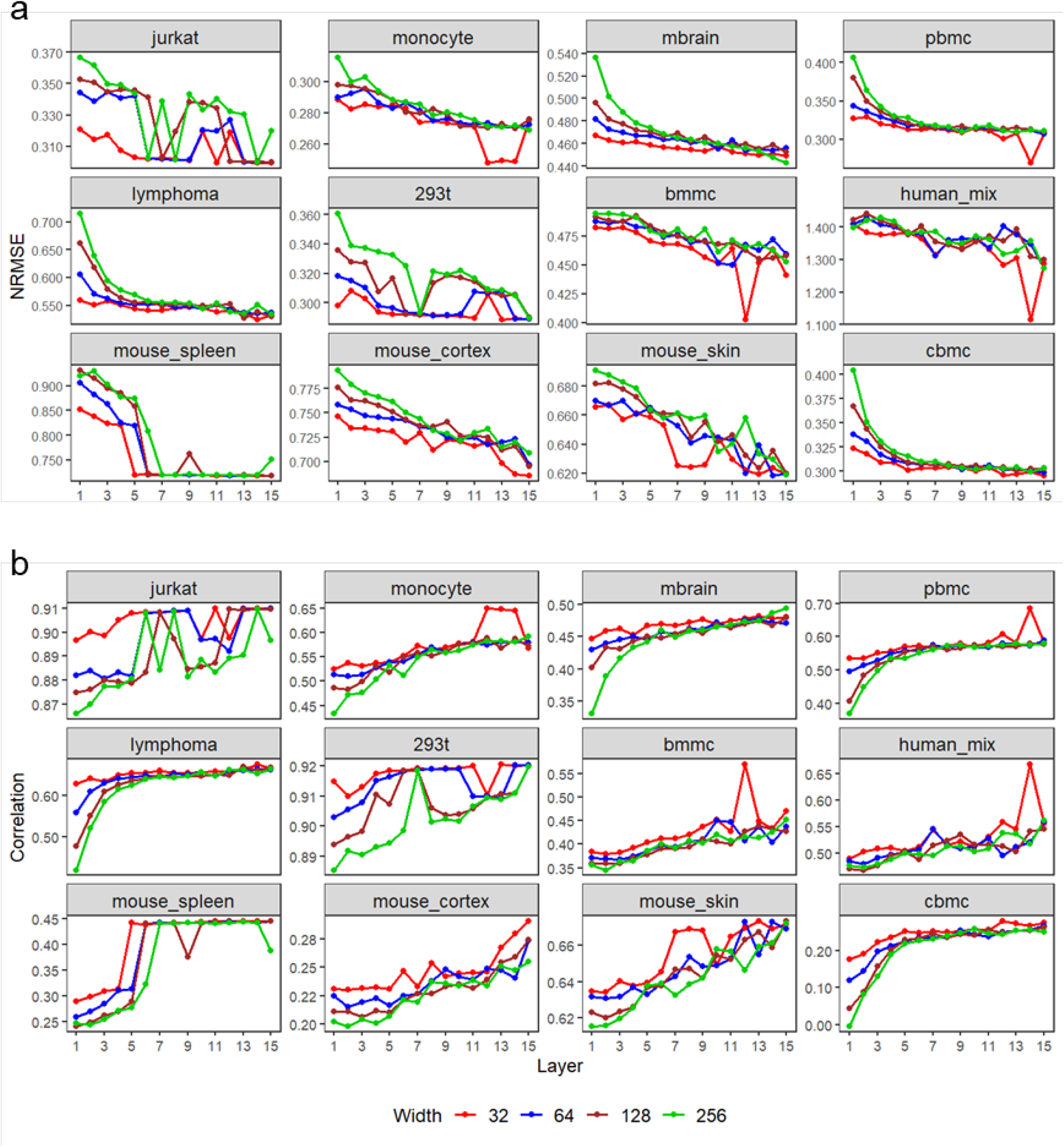
The impacts of autoencoder depth and width on the imputation NRMSE (**a**) and the imputation correlation (**b**) based on the random masking scheme. The number of highly variable genes is set to 3000.

**Supplementary Figure S17.**
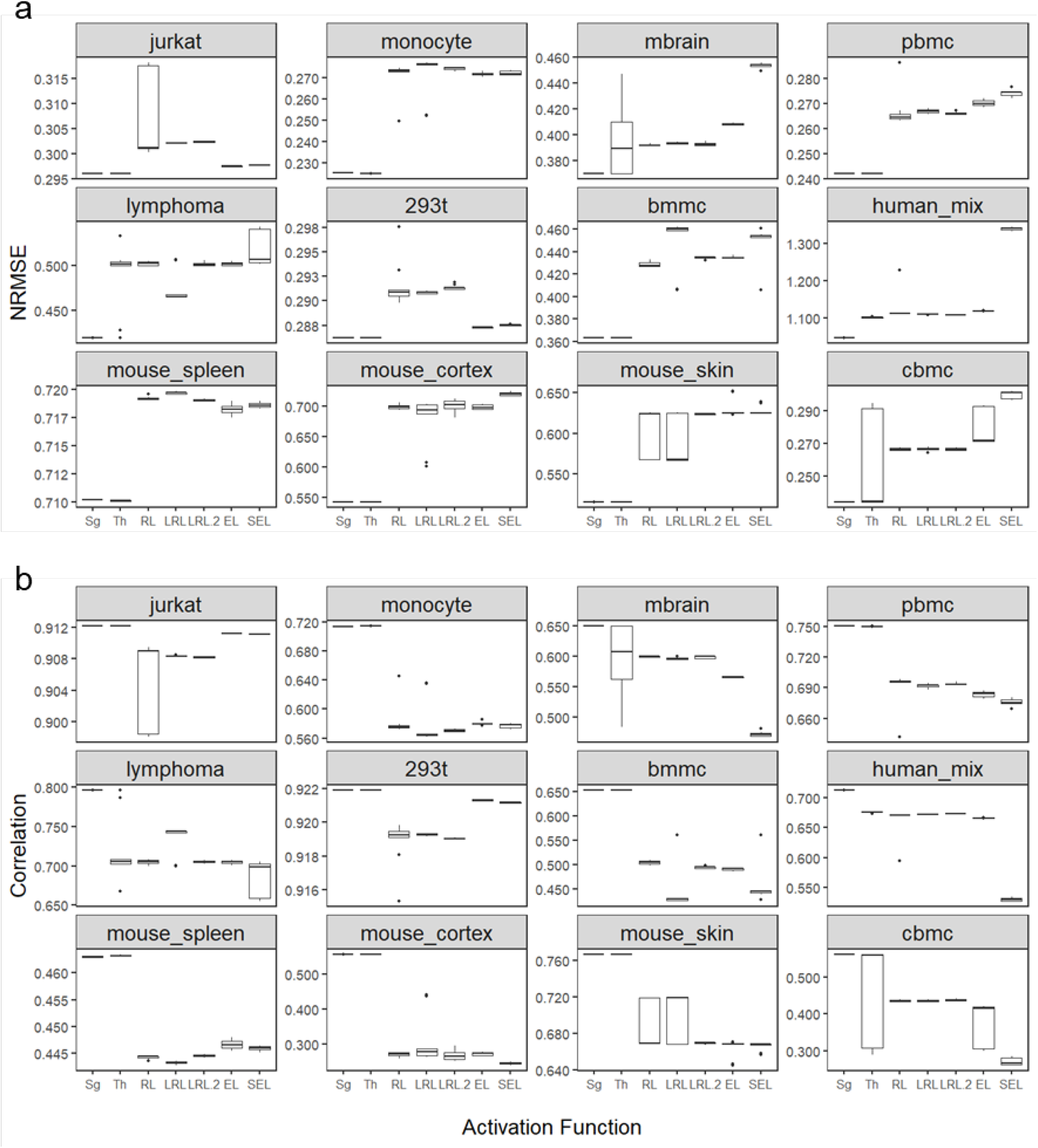
The impact of activation functions on the imputation NRMSE (**a**) and the imputation correlation (**b**) based on the random masking scheme. Sg: sigmoid; Th: tanh; RL: ReLU; LRL: LeakyReLU (=0.01); LRL.2: LeakyReLU (=0.2); EL: ELU; SEL: SELU. The number of highly variable genes is set to 3000.

**Supplementary Figure S18.**
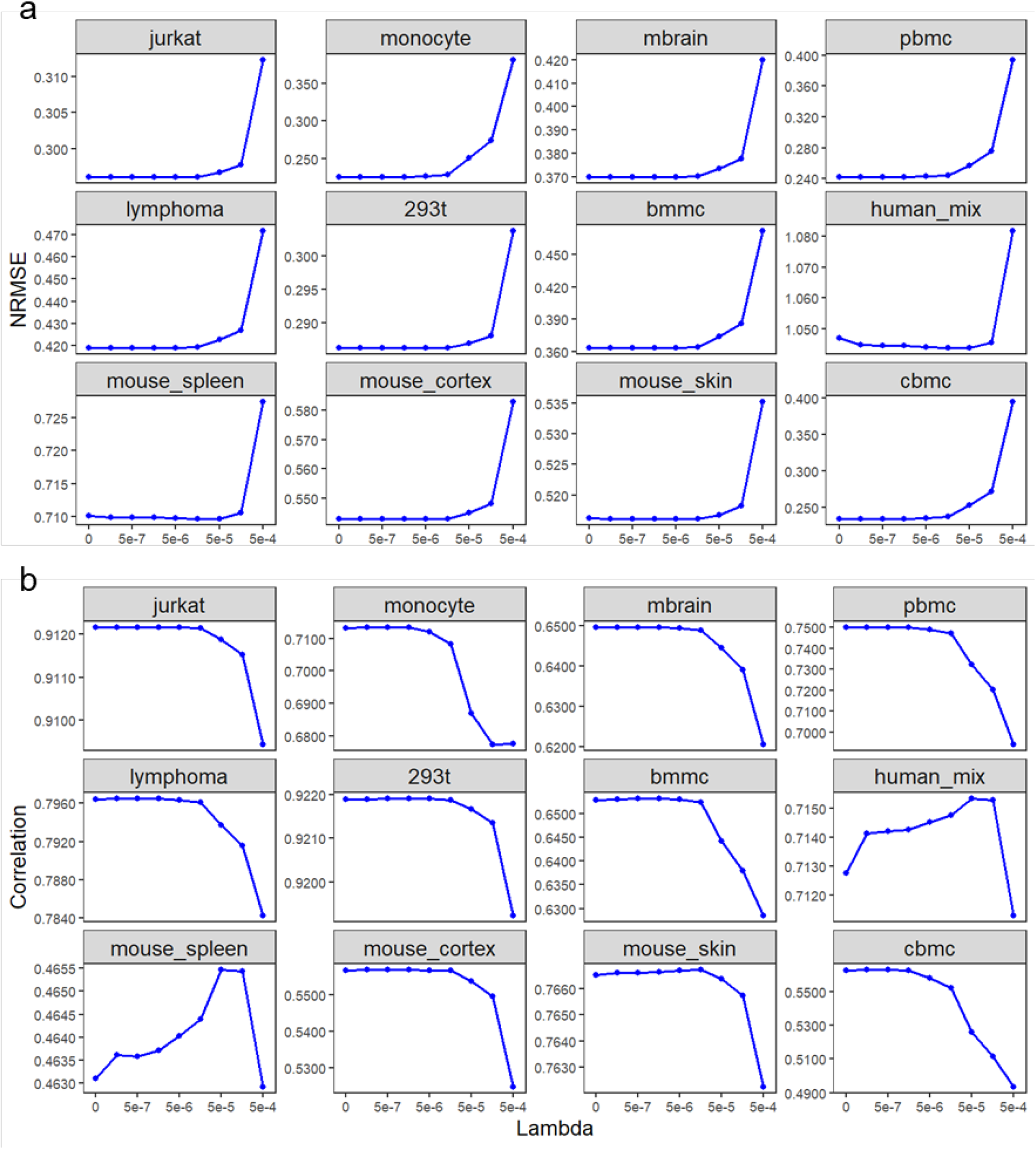
The impact of weight decay regularization on the imputation NRMSE (**a**) and the imputation correlation (**b**) based on the random masking scheme. The number of highly variable genes is set to 3000.

**Supplementary Figure S19.**
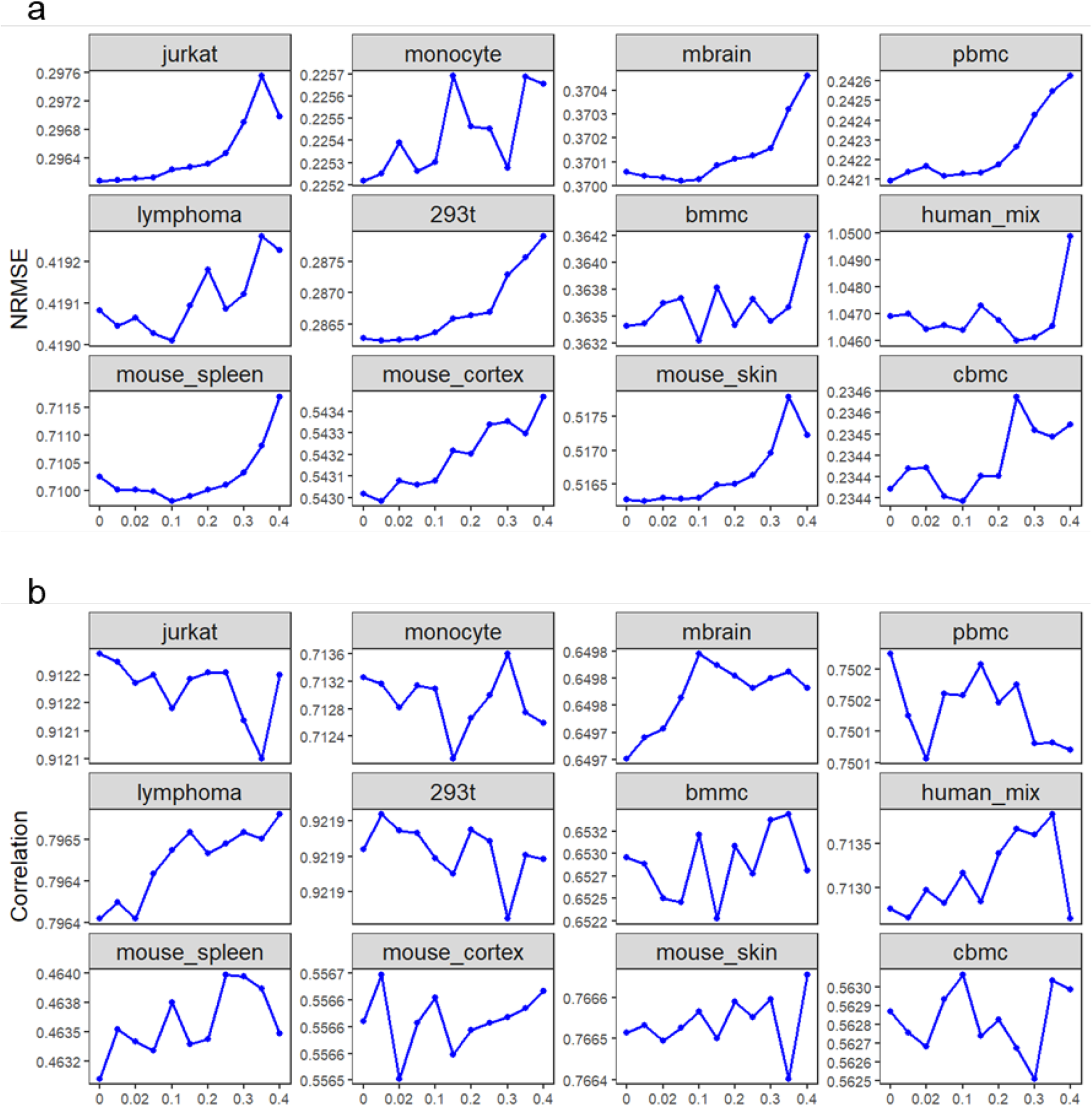
The impact of dropout regularization on the imputation NRMSE (**a**) and the imputation correlation (**b**) based on the random masking scheme. The number of highly variable genes is set to 3000. Some labels on the y-axis are identical due to the low accuracy variability under different dropout rates.

**Supplementary Table S1.** The 12 real scRNA-seq datasets used in the evaluation of overall imputation accuracy.

| Dataset | Tissue/Cell type | Experimental protocol | # of cells | # of genes | Zero rate | Reference |
| --- | --- | --- | --- | --- | --- | --- |
| jurkat | Jurkat cells | 10x Genomics | 3258 | 32738 | 90.23% | 59 |
| monocyte | CD14+ Monocytes | 10x Genomics | 2612 | 32738 | 98.59% | 59 |
| mbrain | Mouse brain cells | 10x Genomics | 9099 | 27998 | 90.97% | 59 |
| pbmc | Peripheral blood mononuclear cells | 10x Genomics | 7783 | 32738 | 97.89% | 59 |
| lymphoma | Lymphoma cells | 10x Genomics | 8412 | 33555 | 96.17% | 59 |
| 293t | 293T Cells | 10x Genomics | 2885 | 32738 | 89.56% | 59 |
| bmmc | Primary bone marrow mononuclear Cells | 10x Genomics | 1985 | 32738 | 97.64% | 59 |
| human_mix | Mixture of HEK293T and MCF7 | Smart-seq-total | 633 | 58660 | 89.58% | 60 |
| mouse_spleen | T-cells from the mouse spleen and small intestine | Smart-seq2 | 574 | 23998 | 84.79% | 61 |
| mouse_cortex | Mouse cortex and hippocampus | Fluidigm C1 | 3005 | 19972 | 81.21% | 62,63 |
| mouse_skin | Mouse skin | Fluidigm C1 | 1422 | 26024 | 90.03% | 64 |
| cbmc | Cord blood mononuclear cells | CITE-seq | 8617 | 14438 | 95.54% | 65 |

**Supplementary Table S2.**
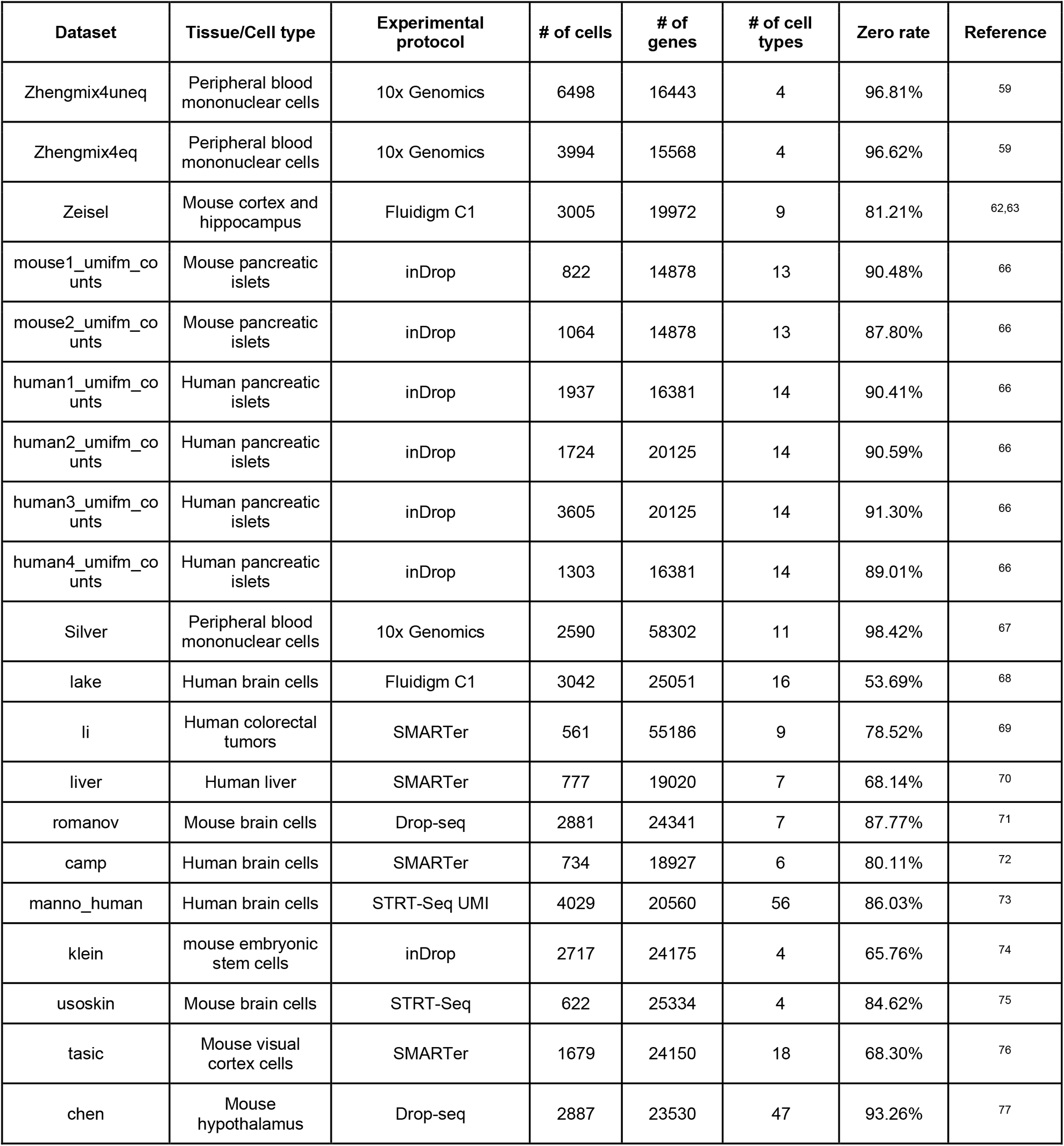
The 20 real scRNA-seq datasets with cell type labels used in the evaluation of cell clustering.

**Supplementary Table S3.**
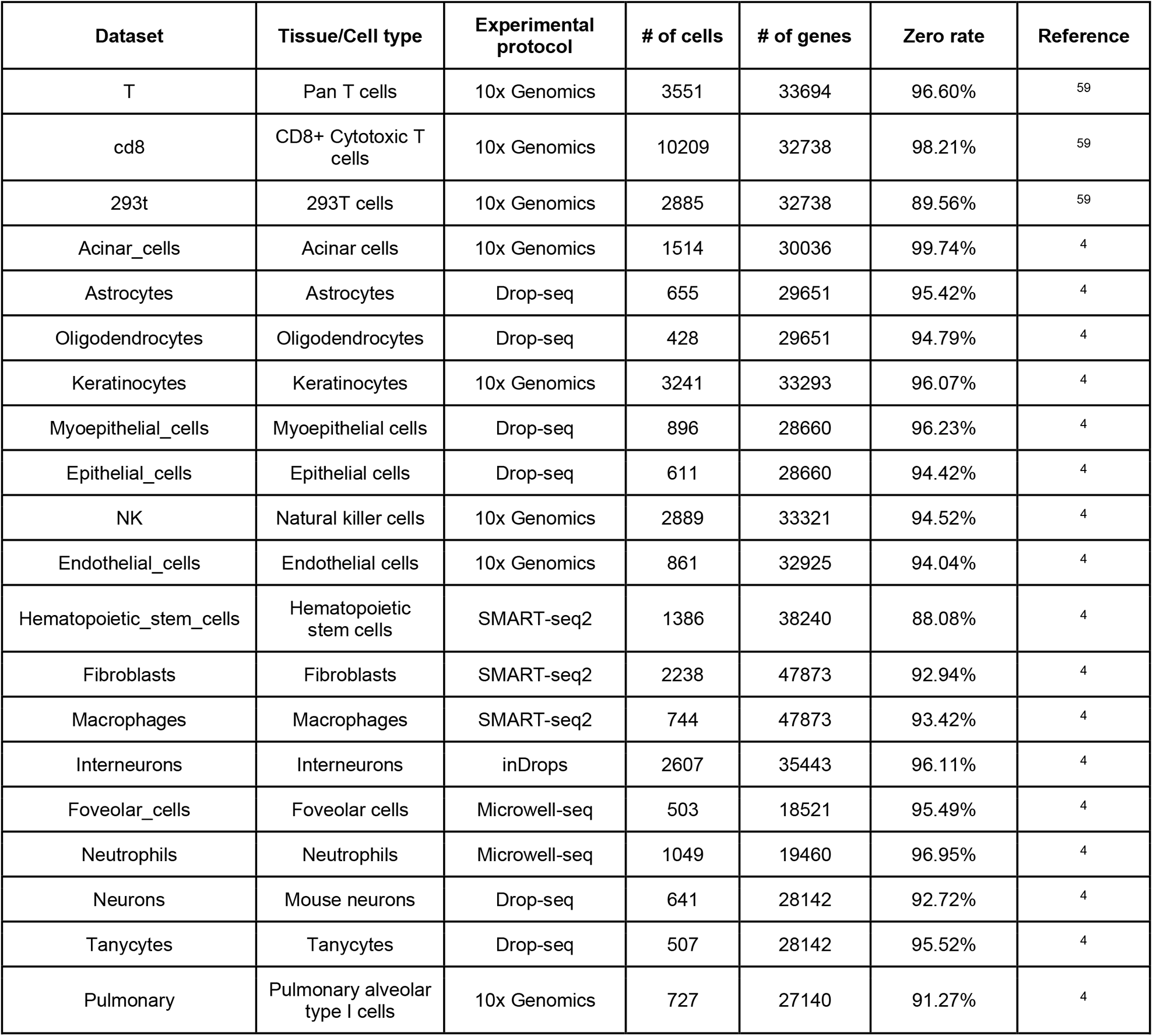
The 20 real scRNA-seq datasets used to generate synthetic datasets in the evaluation of DE gene analysis.

## Reference

1. Hwang, B., Lee, J. H. & Bang, D. Single-cell RNA sequencing technologies and bioinformatics pipelines. Exp. Mol. Med. 50, 96 (2018).

2. Stuart, T. & Satija, R. Integrative single-cell analysis. Nat. Rev. Genet. 20, 257–272 (2019).

3. Chen, G., Ning, B. & Shi, T. Single-Cell RNA-Seq Technologies and Related Computational Data Analysis. Front. Genet. 10, 317 (2019).

4. Franzén, O., Gan, L.-M. & Björkegren, J. L. M. PanglaoDB: a web server for exploration of mouse and human single-cell RNA sequencing data. Database 2019, (2019).

5. Choi, Y. H. & Kim, J. K. Dissecting Cellular Heterogeneity Using Single-Cell RNA Sequencing. Mol. Cells 42, 189–199 (2019).

6. Kiselev, V. Y., Andrews, T. S. & Hemberg, M. Challenges in unsupervised clustering of single-cell RNA-seq data. Nat. Rev. Genet. 20, 273–282 (2019).

7. Van den Berge, K., et al. Trajectory-based differential expression analysis for single-cell sequencing data. Nat. Commun. 11, 1–13 (2020).

8. Lähnemann, D. et al. Eleven grand challenges in single-cell data science. Genome Biol. 21, 31 (2020).

9. Jiang, R., Sun, T., Song, D. & Li, J. J. Statistics or biology: the zero-inflation controversy about scRNA-seq data. Genome Biol. 23, 31 (2022).

10. Bai, Y.-L., Baddoo, M., Flemington, E. K., Nakhoul, H. N. & Liu, Y.-Z. Screen technical noise in single cell RNA sequencing data. Genomics 112, 346–355 (2020).

11. Li, W. V. & Li, J. J. An accurate and robust imputation method scImpute for single-cell RNA-seq data. Nat. Commun. 9, 997 (2018).

12. Azizi, E., Prabhakaran, S., Carr, A. & Pe’er, D. Bayesian Inference for Single-cell Clustering and Imputing. Genomics and Computational Biology vol. 3 46 Preprint at https://doi.org/10.18547/gcb.2017.vol3.iss1.e46 (2017).

13. van Dijk, D. et al. Recovering Gene Interactions from Single-Cell Data Using Data Diffusion. Cell vol. 174 716–729.e27 Preprint at https://doi.org/10.1016/j.cell.2018.05.061 (2018).

14. Gong, W., Kwak, I.-Y., Pota, P., Koyano-Nakagawa, N. & Garry, D. J. DrImpute: imputing dropout events in single cell RNA sequencing data. BMC Bioinformatics 19, 220 (2018).

15. Pierson, E. & Yau, C. ZIFA: Dimensionality reduction for zero-inflated single-cell gene expression analysis. Genome Biol. 16, 241 (2015).

16. Eraslan, G., Simon, L. M., Mircea, M., Mueller, N. S. & Theis, F. J. Single-cell RNA-seq denoising using a deep count autoencoder. Nat. Commun. 10, 390 (2019).

17. Zhang, L. & Zhang, S. Comparison of computational methods for imputing single-cell RNA-sequencing data. IEEE/ACM Trans. Comput. Biol. Bioinform. 17, 376–389 (2020).

18. Hou, W., Ji, Z., Ji, H. & Hicks, S. C. A systematic evaluation of single-cell RNA-sequencing imputation methods. Genome Biol. 21, 218 (2020).

19. Li, W. V. & Li, J. J. A statistical simulator scDesign for rational scRNA-seq experimental design. Bioinformatics 35, i41–i50 (2019).

20. Nair, V. & Hinton, G. E. Rectified linear units improve restricted boltzmann machines. in Icml (2010).

21. Kingma, D. P. & Ba, J. Adam: A Method for Stochastic Optimization. arXiv [cs.LG] (2014).

22. Krizhevsky, A., Sutskever, I. & Hinton, G. E. ImageNet Classification with Deep Convolutional Neural Networks. in Advances in Neural Information Processing Systems (eds. Pereira, F., Burges, C. J. C., Bottou, L. & Weinberger, K. Q.) vol. 25 (Curran Associates, Inc., 2012).

23. Zhao, Z.-Q., Zheng, P., Xu, S.-T. & Wu, X. Object Detection with Deep Learning: A Review. arXiv [cs.CV] (2018).

24. Nwankpa, C., Ijomah, W., Gachagan, A. & Marshall, S. Activation Functions: Comparison of trends in Practice and Research for Deep Learning. arXiv [cs.LG] (2018).

25. Xu, B., Wang, N., Chen, T. & Li, M. Empirical Evaluation of Rectified Activations in Convolutional Network. arXiv [cs.LG] (2015).

26. Clevert, D.-A., Unterthiner, T. & Hochreiter, S. Fast and Accurate Deep Network Learning by Exponential Linear Units (ELUs). arXiv [cs.LG] (2015).

27. Klambauer, G., Unterthiner, T., Mayr, A. & Hochreiter, S. Self-Normalizing Neural Networks. arXiv [cs.LG] (2017).

28. Lu, L., Shin, Y., Su, Y. & Karniadakis, G. E. Dying ReLU and Initialization: Theory and Numerical Examples. arXiv [stat.ML] (2019).

29. Pascanu, R., Mikolov, T. & Bengio, Y. On the difficulty of training Recurrent Neural Networks. arXiv [cs.LG] (2012).

30. Arisdakessian, C., Poirion, O., Yunits, B., Zhu, X. & Garmire, L. X. DeepImpute: an accurate, fast, and scalable deep neural network method to impute single-cell RNA-seq data. Genome Biol. 20, 211 (2019).

31. Talwar, D., Mongia, A., Sengupta, D. & Majumdar, A. AutoImpute: Autoencoder based imputation of single-cell RNA-seq data. Sci. Rep. 8, 16329 (2018).

32. Luecken, M. D. & Theis, F. J. Current best practices in single-cell RNA-seq analysis: a tutorial. Mol. Syst. Biol. 15, (2019).

33. 33. Hao, Y., et al. Integrated analysis of multimodal single-cell data. bioRxiv 2020.10.12.335331 (2020) doi:10.1101/2020.10.12.335331.

34. Finak, G. et al. MAST: a flexible statistical framework for assessing transcriptional changes and characterizing heterogeneity in single-cell RNA sequencing data. Genome Biol. 16, 278 (2015).

35. Tang, F. et al. mRNA-Seq whole-transcriptome analysis of a single cell. Nat. Methods 6, 377–382 (2009).

36. Zappia, L., Phipson, B. & Oshlack, A. Exploring the single-cell RNA-seq analysis landscape with the scRNA-tools database. PLoS Comput. Biol. 14, e1006245 (2018).

37. Fan, L., Zhang, F., Fan, H. & Zhang, C. Brief review of image denoising techniques. Visual Computing for Industry, Biomedicine, and Art 2, 1–12 (2019).

38. Lopez, R., Regier, J., Cole, M. B., Jordan, M. I. & Yosef, N. Deep generative modeling for single-cell transcriptomics. Nat. Methods 15, 1053–1058 (2018).

39. Lin, C., Jain, S., Kim, H. & Bar-Joseph, Z. Using neural networks for reducing the dimensions of single-cell RNA-Seq data. Nucleic Acids Res. 45, e156–e156 (2017).

40. Köhler, N. D., Büttner, M. & Theis, F. J. Deep learning does not outperform classical machine learning for cell-type annotation. bioRxiv 653907 (2019) doi:10.1101/653907.

41. Tensors in Image Processing and Computer Vision. (Springer, London, 2009).

42. Deng, Y., Bao, F., Dai, Q., Wu, L. F. & Altschuler, S. J. Scalable analysis of cell-type composition from single-cell transcriptomics using deep recurrent learning. Nat. Methods 16, 311–314 (2019).

43. Wang, L., Xiao, Q. & Xu, H. Optimal maximin $L_{1}$-distance Latin hypercube designs based on good lattice point designs. aos 46, 3741–3766 (2018).

44. Wang, L., Sun, F., Lin, D. K. J. & Liu, M. Q. Construction of orthogonal symmetric Latin hypercube designs. Stat. Sin. (2018).

45. Wang, L. & Xu, H. A class of multilevel nonregular designs for studying quantitative factors. Stat. Sin. (2022) doi:10.5705/ss.202020.0223.

46. Wang, L., Xu, H. & Liu, M.-Q. Fractional factorial designs for Fourier-cosine models. Metrika (2022) doi:10.1007/s00184-022-00881-2.

47. Goodfellow, I., Bengio, Y., Courville, A. & Bengio, Y. Deep learning. vol. 1 (MIT press Cambridge, 2016).

48. LeCun, Y., Bengio, Y. & Hinton, G. Deep learning. Nature 521, 436–444 (2015).

49. Werbos, P. J. Backpropagation through time: what it does and how to do it. Proc. IEEE 78, 1550–1560 (1990).

50. Kingma, D. P. & Welling, M. Auto-Encoding Variational Bayes. arXiv [stat.ML] (2013).

51. Badsha, M. B. et al. Imputation of single-cell gene expression with an autoencoder neural network. Quantitative Biology 8, 78–94 (2020).

52. Hastie, T., Tibshirani, R. & Friedman, J. The Elements of Statistical Learning: Data Mining, Inference, and Prediction, Second Edition. (Springer Science & Business Media, 2009).

53. Srivastava, N., Hinton, G., Krizhevsky, A., Sutskever, I. & Salakhutdinov, R. Dropout: a simple way to prevent neural networks from overfitting. J. Mach. Learn. Res. 15, 1929–1958 (2014).

54. Hicks, S. C., Townes, F. W., Teng, M. & Irizarry, R. A. Missing data and technical variability in single-cell RNA-sequencing experiments. Biostatistics 19, 562–578 (2018).

55. Paszke, A. et al. PyTorch: An Imperative Style, High-Performance Deep Learning Library. in Advances in Neural Information Processing Systems (eds. Wallach, H., et al.) vol. 32 (Curran Associates, Inc., 2019).

56. Abadi, M., et al. TensorFlow: Large-Scale Machine Learning on Heterogeneous Distributed Systems. arXiv [cs.DC] (2016).

57. Fränti, P. & Sieranoja, S. How much can k-means be improved by using better initialization and repeats? Pattern Recognit. 93, 95–112 (2019).

58. Reza, F. M. An Introduction to Information Theory. (Courier Corporation, 1994).

59. Zheng, G. X. Y. et al. Massively parallel digital transcriptional profiling of single cells. Nat. Commun. 8, 14049 (2017).

60. Isakova, A., Neff, N. & Quake, S. Single cell profiling of total RNA using Smart-seq-total. bioRxiv (2020).

61. Brockmann, L. et al. Molecular and functional heterogeneity of IL-10-producing CD4+ T cells. Nat. Commun. 9, 5457 (2018).

62. Lake, B. B. et al. A comparative strategy for single-nucleus and single-cell transcriptomes confirms accuracy in predicted cell-type expression from nuclear RNA. Sci. Rep. 7, 1–8 (2017).

63. Zeisel, A. et al. Cell types in the mouse cortex and hippocampus revealed by single-cell RNA-seq. Science 347, 1138–1142 (2015).

64. Joost, S. et al. Single-Cell Transcriptomics Reveals that Differentiation and Spatial Signatures Shape Epidermal and Hair Follicle Heterogeneity. Cell Syst 3, 221–237.e9 (2016).

65. Stoeckius, M. et al. Simultaneous epitope and transcriptome measurement in single cells. Nat. Methods 14, 865–868 (2017).

66. Baron, M. et al. A Single-Cell Transcriptomic Map of the Human and Mouse Pancreas Reveals Inter- and Intra-cell Population Structure. Cell Syst 3, 346–360.e4 (2016).

67. Freytag, S., Tian, L., Lönnstedt, I., Ng, M. & Bahlo, M. Comparison of clustering tools in R for medium-sized 10x Genomics single-cell RNA-sequencing data. F1000Res. 7, 1297 (2018).

68. Lake, B. B. et al. Neuronal subtypes and diversity revealed by single-nucleus RNA sequencing of the human brain. Science 352, 1586–1590 (2016).

69. Li, H. et al. Reference component analysis of single-cell transcriptomes elucidates cellular heterogeneity in human colorectal tumors. Nat. Genet. 49, 708–718 (2017).

70. Camp, J. G. et al. Multilineage communication regulates human liver bud development from pluripotency. Nature 546, 533–538 (2017).

71. Romanov, R. A. et al. Molecular interrogation of hypothalamic organization reveals distinct dopamine neuronal subtypes. Nat. Neurosci. 20, 176–188 (2016).

72. Gray Camp, J., et al. Human cerebral organoids recapitulate gene expression programs of fetal neocortex development. Proc. Natl. Acad. Sci. U. S. A. 112, 15672–15677 (2015).

73. La Manno, G. et al. Molecular Diversity of Midbrain Development in Mouse, Human, and Stem Cells. Cell 167, 566–580.e19 (2016).

74. Klein, A. M. et al. Droplet Barcoding for Single-Cell Transcriptomics Applied to Embryonic Stem Cells. Cell 161, 1187–1201 (2015).

75. Usoskin, D. et al. Unbiased classification of sensory neuron types by large-scale single-cell RNA sequencing. Nat. Neurosci. 18, 145–153 (2015).

76. Tasic, B. et al. Adult mouse cortical cell taxonomy revealed by single cell transcriptomics. Nat. Neurosci. 19, 335–346 (2016).

77. Chen, R., Wu, X., Jiang, L. & Zhang, Y. Single-Cell RNA-Seq Reveals Hypothalamic Cell Diversity. Cell Rep. 18, 3227–3241 (2017).

